# Consistency and individual variation in common song types of an urban bird: a multi-population comparison over a decade

**DOI:** 10.64898/2026.09.01.748382

**Authors:** Pablo A. Sosa Negrón, J. Roberto Sosa-López, Gilbert Barrantes, Luis Sandoval

## Abstract

Individual variation in singing behavior is influenced by cultural evolution, environmental constraints, and population dynamics. In closed-ended songbird species, aging may be associated to temporal song variations in individual birds due to vocal apparatus deterioration. However, song variations may also arise as a result of extrinsic factors that induce behavioral changes. In this study, we aimed to examine song-type consistency over 13 years in four populations of the White-eared Ground-sparrow (*Melozone leucotis*, Aves: Passerellidae) along an urban-rural gradient. We also compared within and between-individual variation to determine potential population-level responses. We used long-term data from singing males to quantify five fine structural characteristics (minimum frequency and maximum frequencies, bandwidth, peak frequency, and song duration) for two most common song types from each population. Song consistency differed between fine-structure characteristics, song types, and populations. Contrary to our expectations, we found that the Highly urbanized population had greater between-individual variance, associated with anthropogenic disturbances such as habitat structure and noise pollution, suggesting non-uniform individual responses. The Medium, Low urbanized, and Rural populations, on the other hand, had higher levels of within-individual variance, associated to individuals’ age. Our study brings insight into how both intrinsic and extrinsic factors can influence song consistency and highlights how cultural evolution within a species can diverge across populations experiencing distinct environmental contexts.

## INTRODUCTION

Animals produce acoustic signals in non-sexual (i.e., signaling food, predator, or location) and sexual contexts (i.e., competing for mates and territories) (Searcy & Andersson 1986; Dreher & Pröhl 2014; Liu et al. 2021). Consequently, acoustic signals are under constant selective pressures (Podos et al. 2004; Sierro et al. 2023). In the case of birds, males may announce to other males their dominance or may indicate females their physical condition using vocalizations (Kroodsma & Byers 1991). This is especially relevant for social interactions with conspecifics to secure their reproductive success and survival (Warrington et al. 2014; Keen et al. 2016). Furthermore, communicating between conspecifics is facilitated by individual distinctiveness, which are signals that differentiate traits associated with their dominance status, health, or size between individulas (Gil & Gar 2002; Tibbets & Dale 2007; Schmidt et al. 2014).

Vocalizations in song learning bird species may vary on three scales (Podos & Warren 2007; Medina & Francis 2012). First, differences between populations, in the form of dialects, arise in geographically separated populations within distinct environmental and social contexts (Wilkins et al. 2013) and are maintained if they are effectively communicating within the local environment (Luther & Baptista 2011; Graham et al. 2017). Second, individuals vary from each other because of errors or improvisations in songs acquired during their learning phase, or because of physical features that are intrinsic to the individuals (Wilson & Mennill 2010; Mennill 2011). Third, variations within individuals occur in terms of song consistency, the ability to produce songs, syllables, or elements with limited spectral variation over time, and have been previously related to male quality in certain species (Taff & Freeman-Gallant 2016). Song variations within individuals can occur across both short temporal scales (i.e., within the same year or breeding season) and longer temporal scales (i.e., between years) (Derryberry 2011; Vargas-Castro et al. 2015).

Bird songs often contain stable acoustic features that facilitate social communication and individual recognition, yet son structure can also change through time because of cultural evolution, environmental influences, and age-related processes (Wiley 2013; Derryberry 2011; Vargas-Castro et al. 2015). At the population level, song traditions and dialects may persist for decades despite continuous individual turnover, reflecting the long-term transmission of culturally inherited vocal traits (Planqué et al. 2014). At the individual level, maintaining highly stereotyped song characteristics has been proposed to reflect aspects of quality because producing repeatable vocal performances may require precise motor control and coordination (Narango & Rodewald 2018). Further, subtle individual variation can arise through environmentally induced plasticity when adjustments provide greater benefits than costs (Gross et al. 2010), or through age-related processes such as ontogenetic changes and deterioration of the vocal apparatus (Rivera-Gutierrez et al. 2012).

Studying between and within-individual variation in behavior has gained interest in behavioral ecology as it can provide insight on microevolutionary processes, population dynamics, and how animals may respond to their environmental contexts (Bolnick et al. 2011; Westneat et al. 2015). In birds, song works as an honest signal of a male’s body condition because it reflects factors such as aggressiveness, intensity of intersexual competition, and resource availability (Szymkowiak & Kuczyński 2017). Consequently, within-individual variation can also indicate the underlying mechanisms behind plasticity and consistency (Nussey et al. 2007; Martin et al. 2017). However, plasticity can also be the result of adaptive responses, such as morphological or physiological constraints caused by aging or environmental changes that may alter trait expression (Gotthard & Nylin 1995; Friis et al. 2022). Whichever the case, song variation or consistency within-individuals may indicate how bird populations respond to changing environments.

Individual song consistency can be used to determine how song variation in birds inhabiting under distinct environmental conditions is associated to dialect evolution within and between populations. For this study, we aimed to examine individual male song type consistency over 13 years in an urban surviving species, the White-eared Ground-Sparrow *(Melozone leucotis*, Aves: Passerellidae). This species exhibits individually distinctive traits in its male solo songs, and it has been demonstrated that males maintain certain stability in their structure between short time periods (Sandoval et al. 2014; Bonilla-Badilla 2021). They are year-round territorial species and males produce their solo songs primarily for territorial defense and mate attraction (Sandoval & Mennill 2012; Sandoval et al. 2013, 2016).

White-eared Ground-Sparrows present microgeographic variations in their songs (dialects), with populations presenting different song types (Sandoval et al. 2014; Bonilla-Badilla 2021). For our study, we used four populations within a rural-urban gradient, which allowed us to compare how male solo songs may vary over time depending on their environmental context. We compared within and between-individual solo song variation, to quantify how much singing males express song consistency on solo songs over time and between habitats (between and within populations). Although song consistency may be important to maintain social status, we predict that within-individual variance in common song types from males will increase over time, because males are older and singing organs age, leading to a decrease in consistency over time. Specifically, we expect that within-individual variation in spectro-temporal characteristics of solo songs, such as minimum frequency, maximum frequency, bandwidth, peak frequency, and song duration, will increase over time. This study allows also to examine the course of cultural evolution in different populations of the same species that potentially face varying degrees of environmental disturbances.

## METHODS

### Ethical note

All procedures involving animals were conducted in accordance with the ethical standards for the use of animals in research. The study protocol was reviewed and approved by the Institutional Committee for the Care and Use of Animals (CICUA) of the University of Costa Rica and by the Research Committee of the School of Biology, University of Costa Rica.

Capture, handling, and sampling of wild birds were carried out under permits issued by the *Sistema Nacional de Áreas de Conservación* (SINAC), Costa Rica. Every effort was made to minimize handling time, stress, and disturbance to the animals throughout the study.

### Study sites

We recorded White-eared Ground-Sparrows in four populations within the Costa Rican Central Valley (Fig 1) during 14 consecutive breeding seasons from 2011 to 2024: (1) Universidad de Costa Rica, Rodrigo Facio Campus in San Pedro (Higly urbanized: 09°56’N, 84°03’W, 1200m), composed of secondary forest growth, fragmented green patches, gardens, and volume large number of buildings (Juarez et al. 2020). (2) Instalaciones Deportivas, Universidad de Costa Rica in San Pedro (Medium urbanized: 9°56’N, 84° 03’W, 1200m), which is a mix of secondary forest growth, gardens, and few buildings (Juarez et al. 2020). (3) Jardín Botánico Lankester, Cartago (Low urbanized: 9°50’N, 83°53’W, 1370m), an area composed of secondary growth forest, gardens, and sparsely buildings (Juarez et al. 2020). (4) Getsemaní, Heredia (Rural: 10°01’N, 84°05’W, 1350m), primarily composed of coffee plantations, fragmented thickets and secondary growth forest (Sandoval et al., 2015).

**Figure 1.**
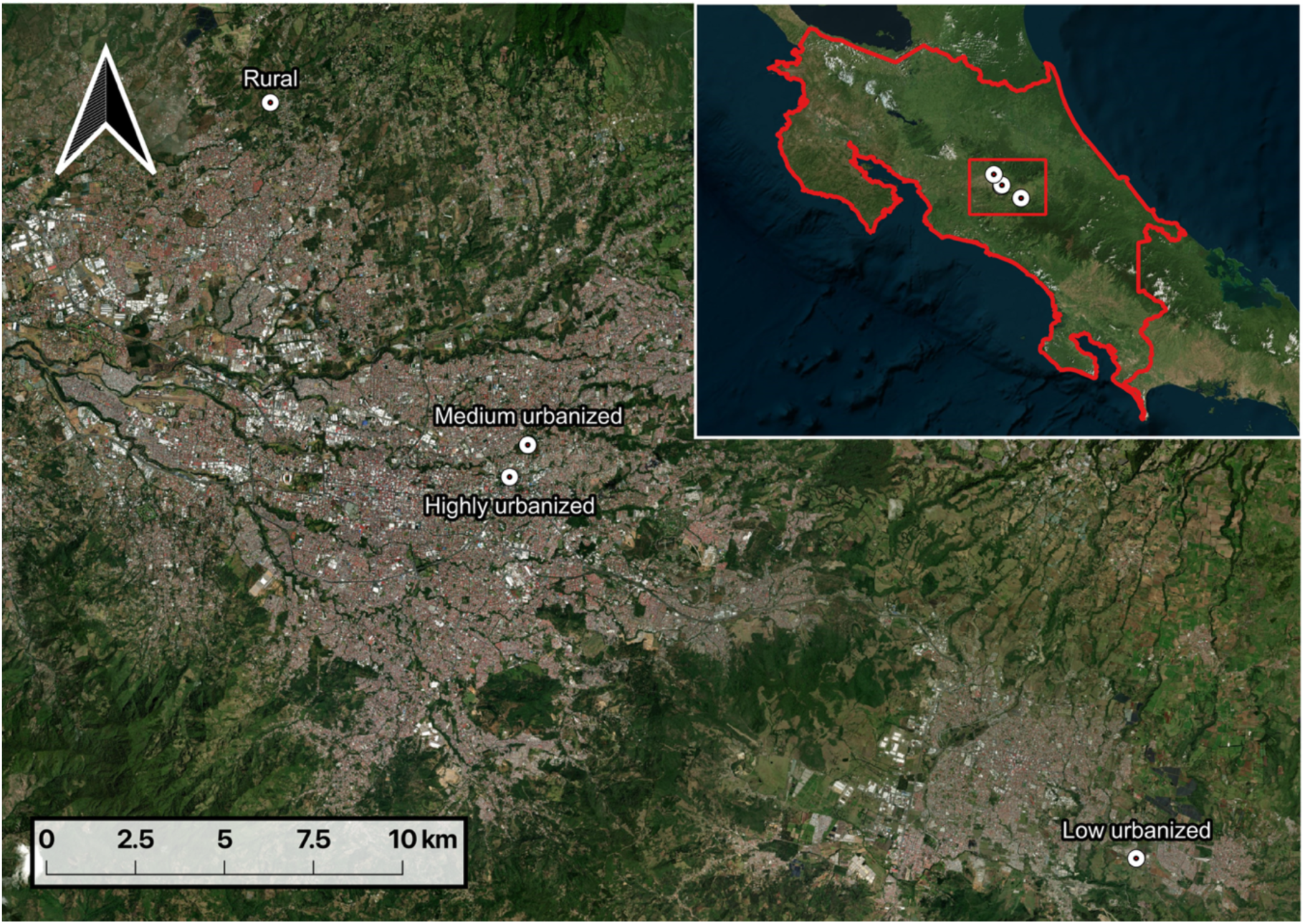
Location of the four populations of White-eared Ground-sparrow males recorded within the Costa Rican Central Valley.

### Song recordings

We recorded White-eared Ground-Sparrow male solo songs between April and June from 2011 through 2024, at the onset of the species’ breeding season (Sandoval & Mennill 2012).

Recordings were conducted between 0500 and 0600 h, when this species is most vocally active (Sandoval et al. 2016), using the focal recording method (Sandoval et al. 2024) with Sennheiser ME66 shotgun microphones and Marantz PMD661digital recorders (recording format: WAVE; sampling rate: 44.1 kHz; 16-bit accuracy). Males were previously banded with a unique colored ring combination and a metallic number ring. This allowed us to identify and record them during consecutive years. We used information of 40 males (10 from each population) that were recorded two or more years to analyze if their common song types varied significantly during these 13 years. We defined common song types as those songs that were produced by at least 68% of males within a population and persist for consecutive years (Bonilla-Badilla 2021). These songs vary according to each population’s dialect (Sandoval et al. 2014; Bonilla-Badilla 2021). We selected males that sang for at least 2 years and that had at least 10 songs recorded from each year.

### Song classification and measurements

We classified songs based on their syntactic structure observed on spectrograms using Raven Pro 1.4 (Cornell Lab of Ornithology, Ithaca, NY, USA) following Sandoval et al. (2014). To ensure accurate identification of song types, we compared the recorded songs to a song type catalog developed by Sandoval et al. (2014). Song types vary between individuals since each male added or omitted introductory elements or varied the length of the terminal trill.

Nonetheless, we classified as the same song type, all songs that had similar overall fine structural features and contained the same number of elements in the middle section following Sandoval et al. (2014) classification. We measured five fine structure features in each song: 1) minimum frequency (Hz), 2) maximum frequency (Hz), 3) bandwidth (Hz), 4) peak frequency(Hz), and 5) song duration (s). At the end, we selected only the two common song types of each population for analyses since these were common among males over years (Bonilla Badilla 2021). These were: T14 and T18 for the Highly urbanized; T13 and T35 for the Medium urbanized; T21 and T42 for the Low urbanized; and T2 and T3 for the Rural site (Fig 2).

**Figure 2.**
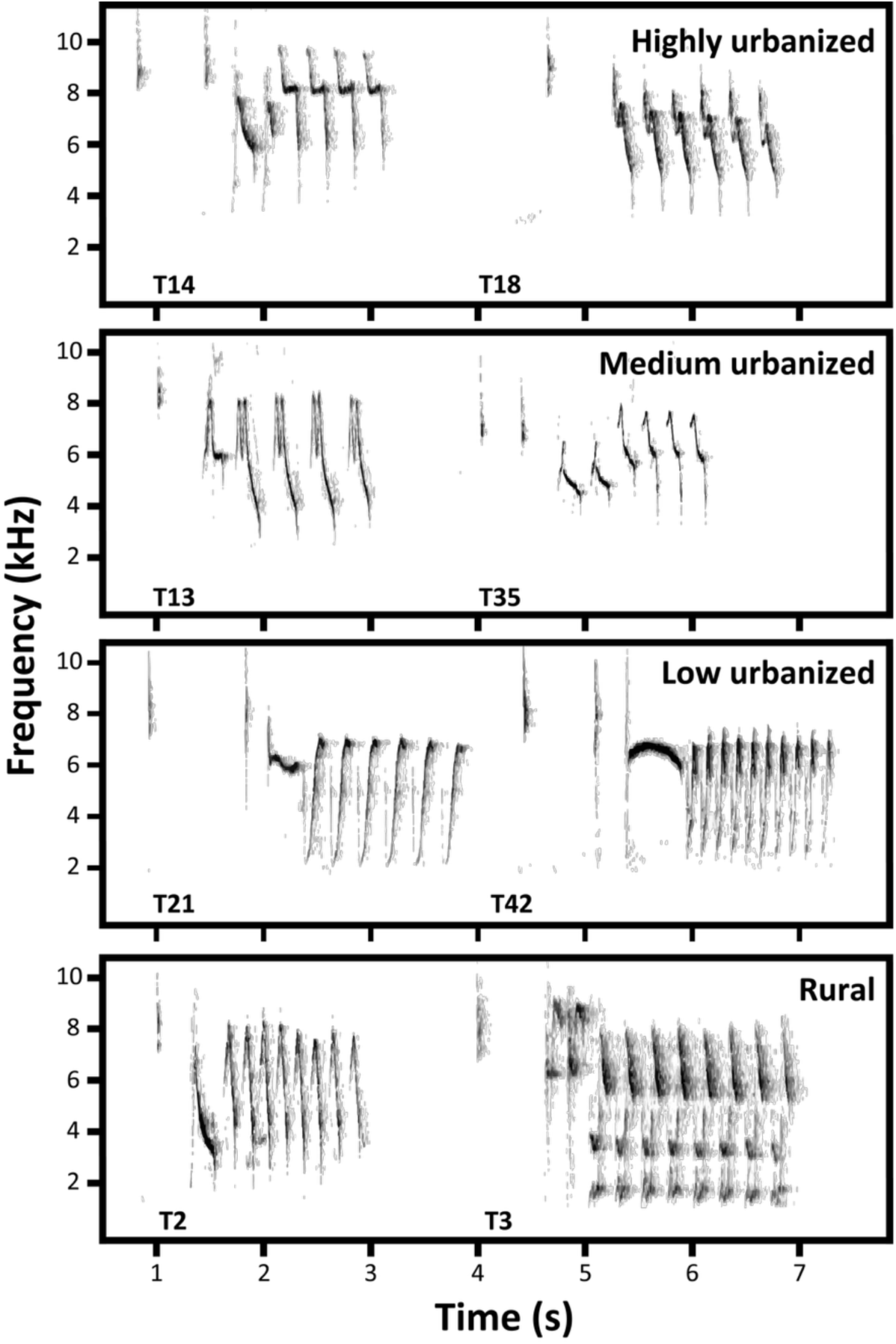
Sonograms of each of the two common song types from each studied population.

### Statistical analysis

All statistical analyses were conducted in R (R Core Team 2026). To determine whether recorded males could be distinguished based on common songs within each population, we performed a multivariate analysis of variance (MANOVA). We used male identity as the independent variable and all fine-structure song features as response variables, excluding DF to reduce collinearity. To quantify the degree of temporal variation (over years) in male songs, we performed linear mixed models (LMM) for each common song type using packages ‘lme4’ to fit models (Bates et al. 2015) and ‘lmerTest’ to extract *p*-values (Kuznetsova et al. 2017). In these models, we included *Year* as a fixed factor, male individuals were added as random factors, and the fine structure features (minimum frequency, maximum frequency, bandwidth, peak frequency, and song duration) as response variables, each in separate models. We analyzed between 10 and 167 songs per year per individual (mean ± SD = 46.1 ± 35.6), across a sampling period of two to four years per individual. We interpreted the models’ variance of the random intercept (*Vind*) as between-individual, and the residual variance *(Ve)* as an approximation to within-individual variance (Van de Pol & Wright 2009; Dingemanse & Dochtermann 2013).

Thus, *Ve* > *Vind* would indicate higher within-individual variance, and *Vind* > *Ve* higher between-individual variance. We acknowledge that *Ve* may also include measurement errors and other unaccounted sources of variation (Westneat et al. 2015). However, potential errors were reduced by including *Year* as a fixed effect, since this way it accounts for variation attributable to temporal trends (Dingemanse & Dochterman 2013). Individual repeatability (R) was calculated as *Vind* / (*Vind* + *Ve*), providing a standardized estimate of between-individual differentiation (Nakagawa & Schielzeth 2010).

By performing separate models for each common song type, we ensured that *Vind* specifically captured individual distinctiveness without the confounding effects of common song type differences. We also performed an ANOVA to determine how distinct males were from one another and a Tukey test to verify where exactly the difference occurred (see Table S1).

Descriptive statistics for all fine-structure features by song type and population are provided in Supplementary Material (Figs S1–S5).

## RESULTS

From the 40 males recorded, 34 males accomplished the requirements for the analyses: 9 males from the Highly urbanized, 8 from the Medium urbanized, 7 from the Low urbanized, and 10 from the Rural site. We processed a total of 118 h of recordings and measured a total of 5,143 songs. There was data available for nearly every year, except for 2020. Because of limitations during to the COVID-19 lockdown, recordings were very reduced and did not meet the requirements to be used in the analysis. We found that males from the four populations exhibited significant individual distinctiveness for each common song type (Table S2), thereby supporting the consistency and accuracy of our visual classification of song types.

We found differences in the between-individual and within-individual variance for the spectro-temporal song characteristics per population (Table 1). Temporal trends were different among fine-structure features and among song types. Song types from the Highly and Medium populations had a mean minimum frequency above 3000 Hz, whereas from the Low urbanized and Rural were mostly between 2000 and 4000 Hz (Fig S1). For the common song T14 of the Highly urbanized population, minimum frequency and song duration increased over time, while maximum frequency and bandwidth decreased (Table 1). For the Highly urbanized population T18 common song, maximum frequency and bandwidth decreased over time, and song duration increased (Table 1). In T13 and T35, both songs from the Medium urbanized population, minimum frequency and song duration increased, while maximum frequency and bandwidth decreased for T13 through the years (Table 1). Minimum frequency decreased temporally in songs from the Rural (T2 and T3) and Low urbanized (T21 and T42) populations (Table 1). Peak frequency decreased in T2 and T35, but it increased in T3, T13, and T21(Table 1). Bandwidth increased in T3 and T42 (Table 1). No types of variance or temporal trends were observed on maximum frequency and song duration from T2, and T21 and T42 (Table 1). Also, bandwidth did not change for T2 and T21, as well as peak frequency in T42 (Table 1).

**Table 1.**
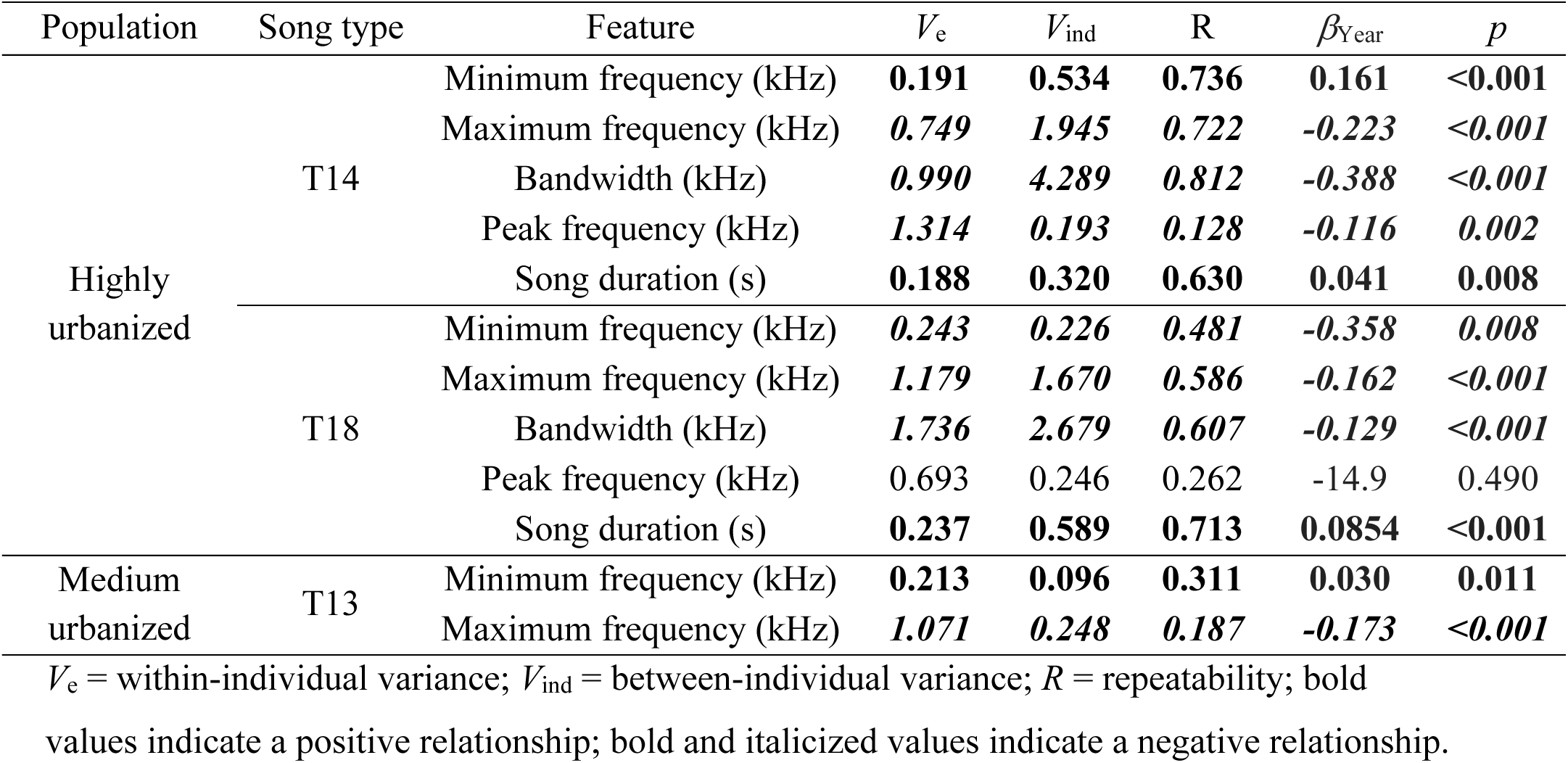
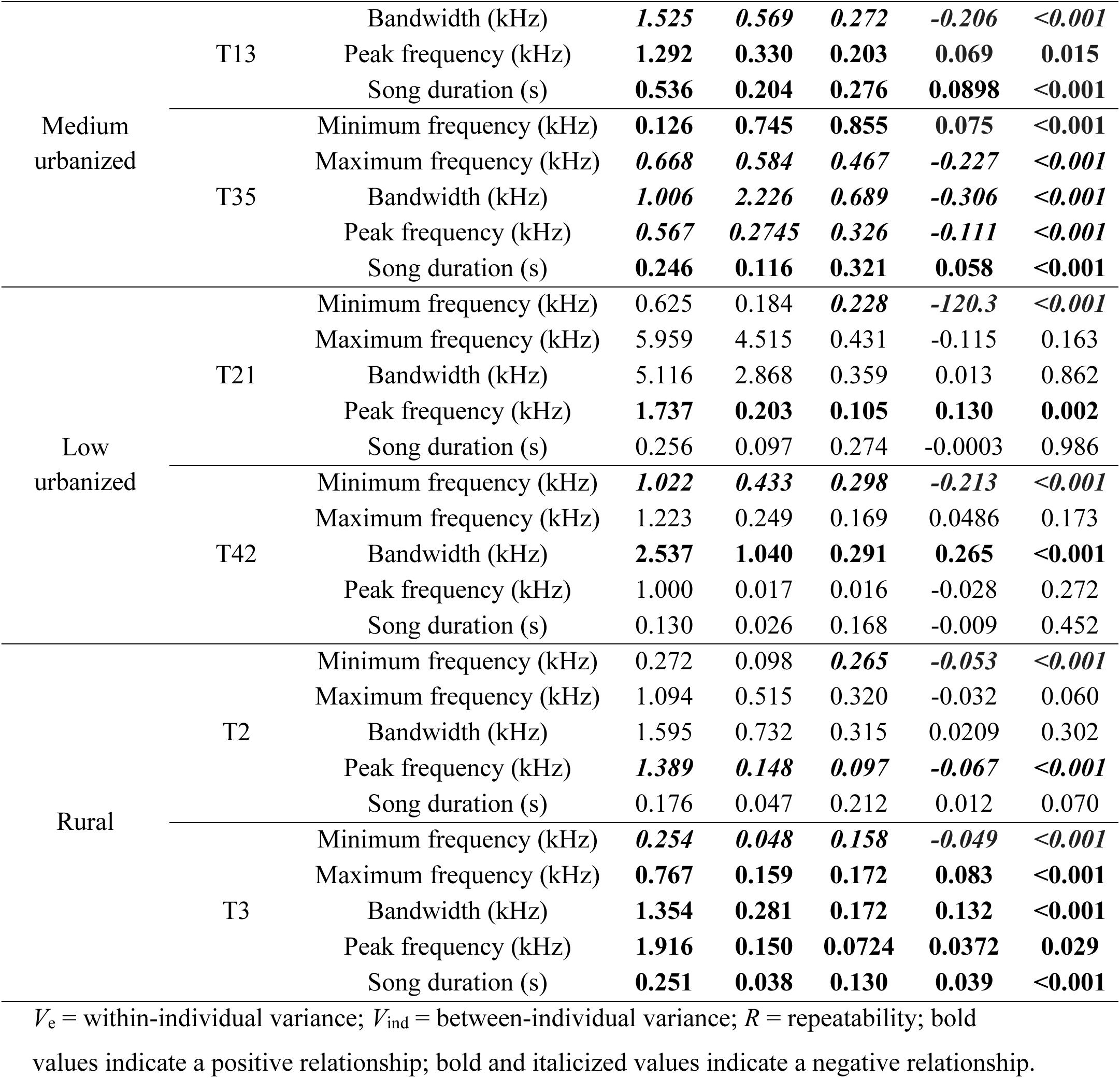
Temporal variance in the fine structure features of common song types of four populations of the White-eared Ground-Sparrow along an urban gradient.

Significant between-individual variation was present in nearly all fine-structure features of both Highly urbanized population’s common song types, whereas in the Medium urbanized, this was only observed for a single feature in one song type (Table 1). For T14 specifically, between-individual variance (*Vind* > *Ve*) was higher for minimum frequency (Table 1; Fig 3A), maximum frequency (Table 1; Fig 4A), bandwidth (Table 1; Fig 5A), and song duration (Table 1; Fig 6A). In T18, between-individual variance was higher for maximum frequency (Table 1; Fig 4A), bandwidth (Table 1; Fig 5A), and song duration (Table 1; Fig 6A). Lastly, in T35 between-individual variance was higher for bandwidth (Table 1; Fig 5B).

**Figure 3.**
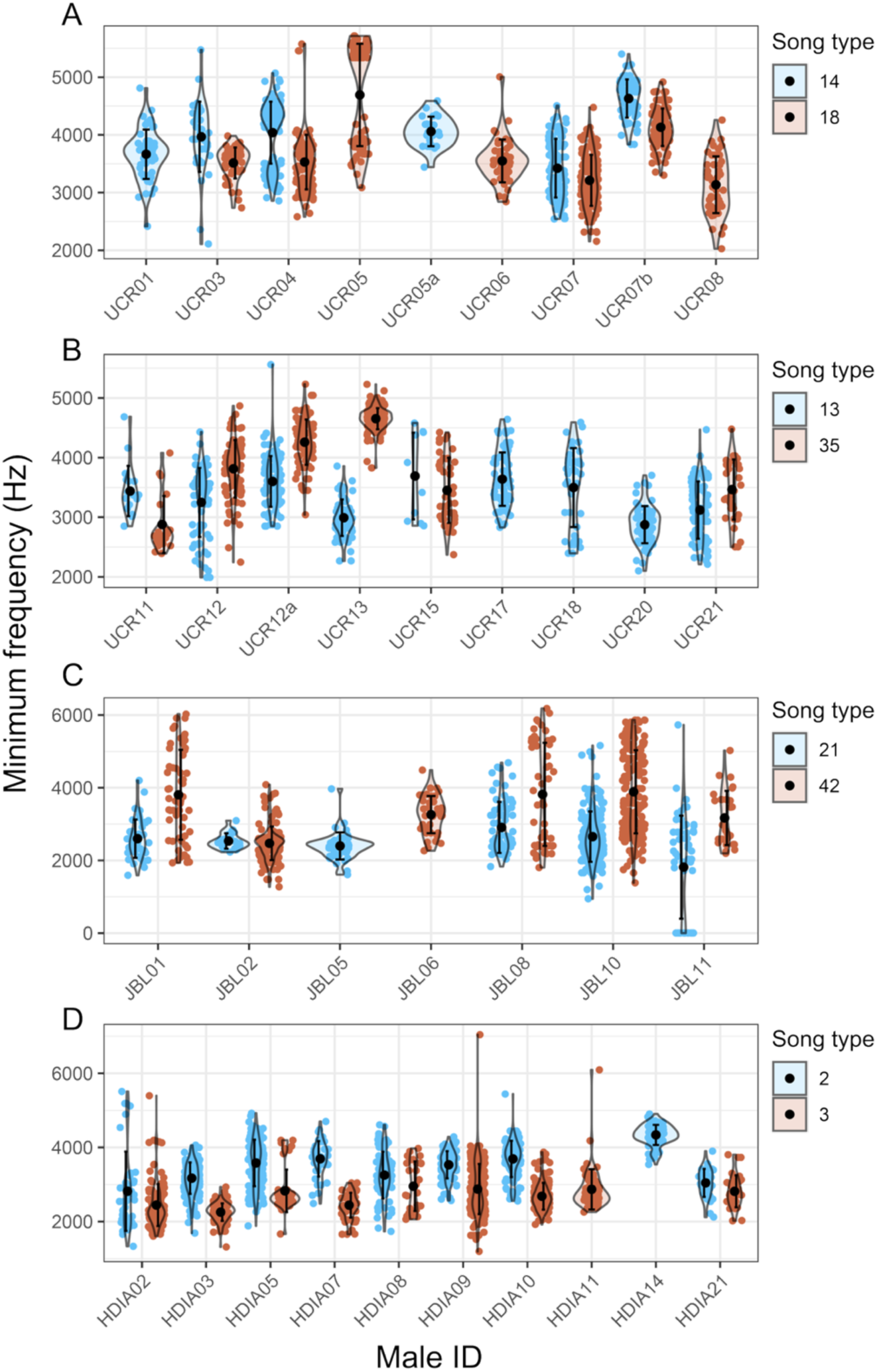
Temporal variance in minimum frequency in common song types from the (A) Highly urban, (B) Medium urban, (C) Low urban, and (D) Rural populations. Central black dots represent mean minimum frequency for each male within the 13-time frame, error bars represent the standard deviation, and colored points represent the measurements of the minimum frequency

**Figure 4.**
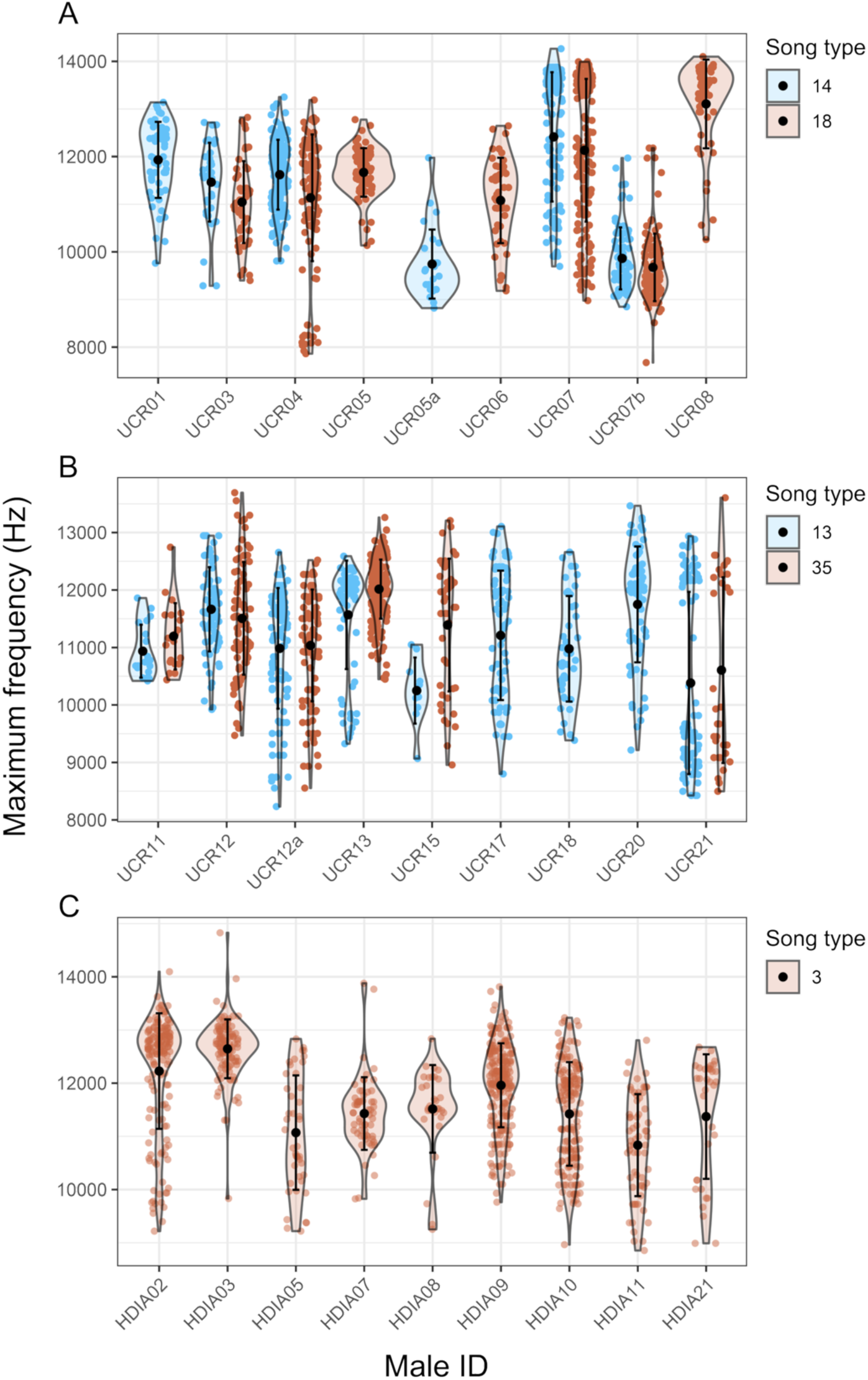
Temporal variance in maximum frequency in common song types from ((A) Highly urban, (B) Medium urban, (C) Low urban, and (D) Rural populations. Central black dots represent mean maximum frequency for each male within the 13-time frame, error bars represent the standard deviation, and colored points represent a measurement.

**Figure 5.**
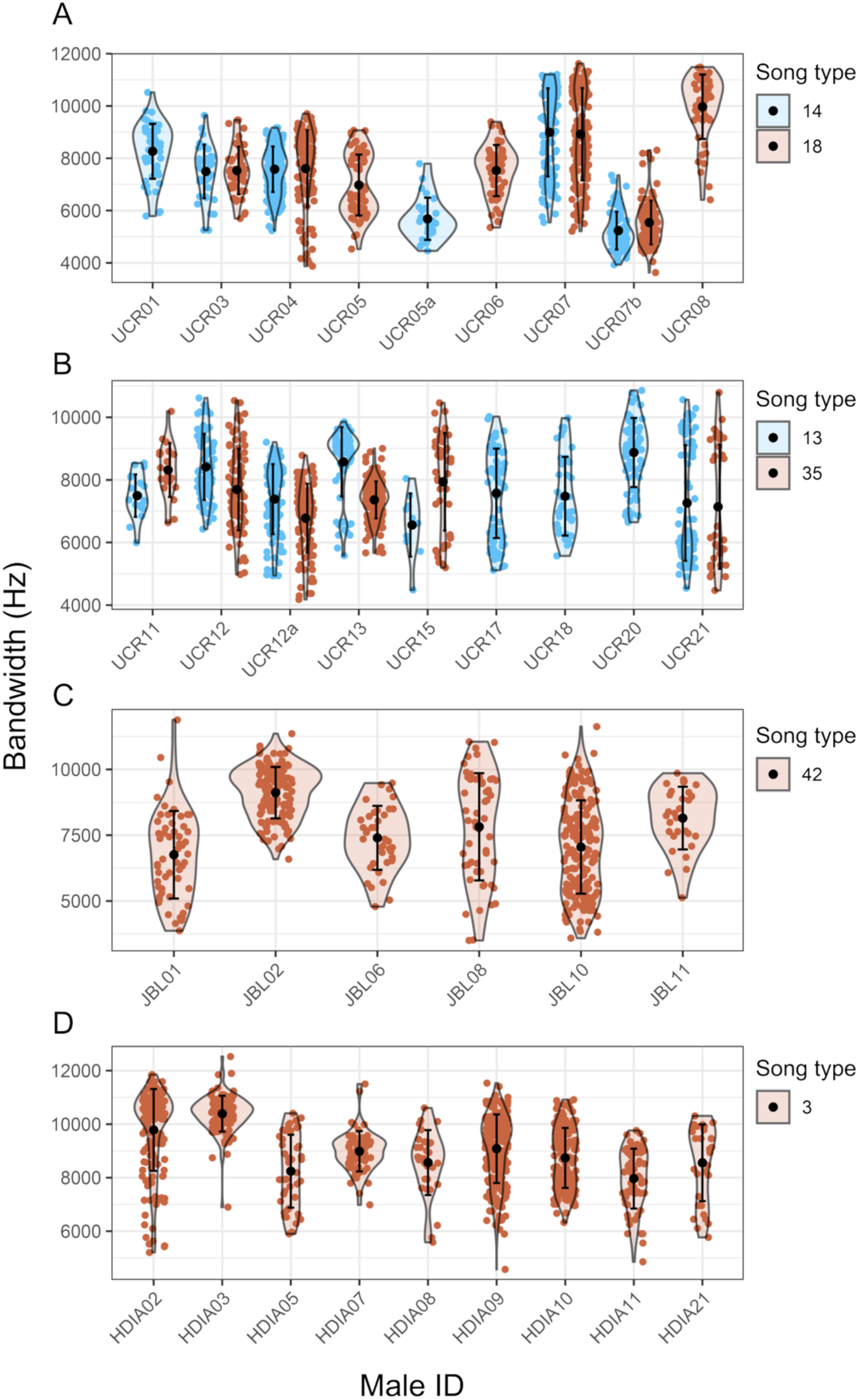
Temporal variance in bandwidth in common song types from (A) Highly urban, (B) Medium urban, (C) Low urban, and (D) Rural populations. Central black dots represent mean frequency range for each male within the 13-time frame, error bars represent the standard deviation, and colored points represent a measurement.

**Figure 6.**
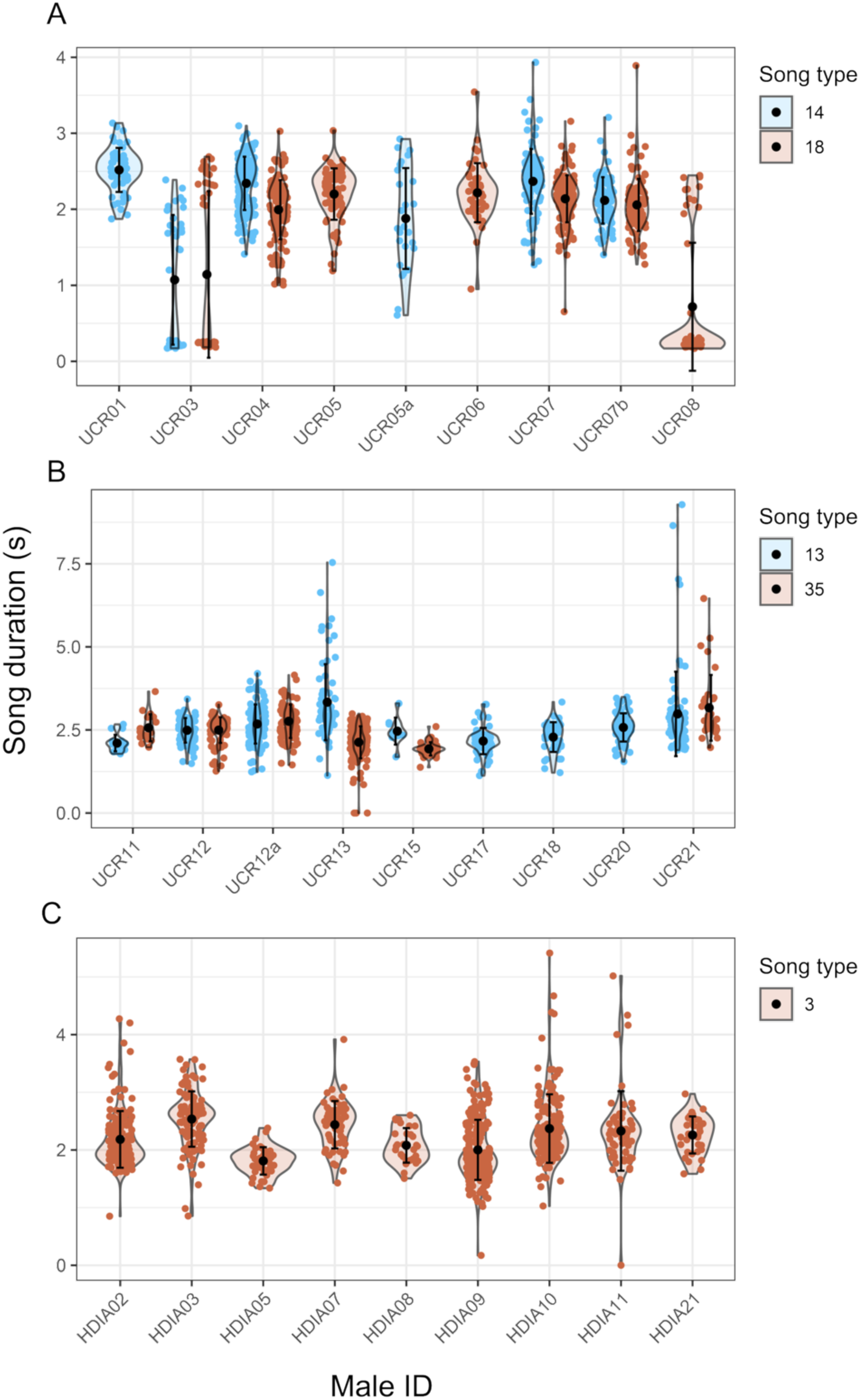
Temporal variance in song duration in common song types from ((A) Highly urban, (B) Medium urban, (C) Low urban, and (D) Rural populations. Central black dots represent mean song duration for each male within the 13-time frame, error bars represent the standard deviation, and colored points represent a measurement.

Within-individual variation was mostly higher (*Vind* < *Ve*) in the Medium, Low urbanized, and Rural populations’ common song types (Table 1). We found higher within-individual variance in minimum frequency in T13 and T35(Table 1; Fig 3B); T21 and T42 (Table 1; Fig 3C); and T2 and T3 (Table 1; Fig 3D). Within-individual variance was also higher in the maximum frequency from T13 and T35 (Table 1; Fig 4B), and T3 (Table 1; Fig 4C). The same pattern was observed in bandwidth from T13 (Table 1; Fig 5B), T42 (Table 1; Fig 5C), and T3 (Table 1; Fig 5D). In peak frequency, we also found higher within-individual variance in T13 and T35 (Table 1; Fig 7B), T21 (Table 1; Fig 7C), and T2 and T3 (Table 1; Fig 7D). Finally, within-individual variance was higher in song duration from T13 and T35 (Table 1; Fig 6B), and T3 (Table 1; Fig 6C). To observe the specific differences between individuals, refer to Table S2.

**Figure 7.**
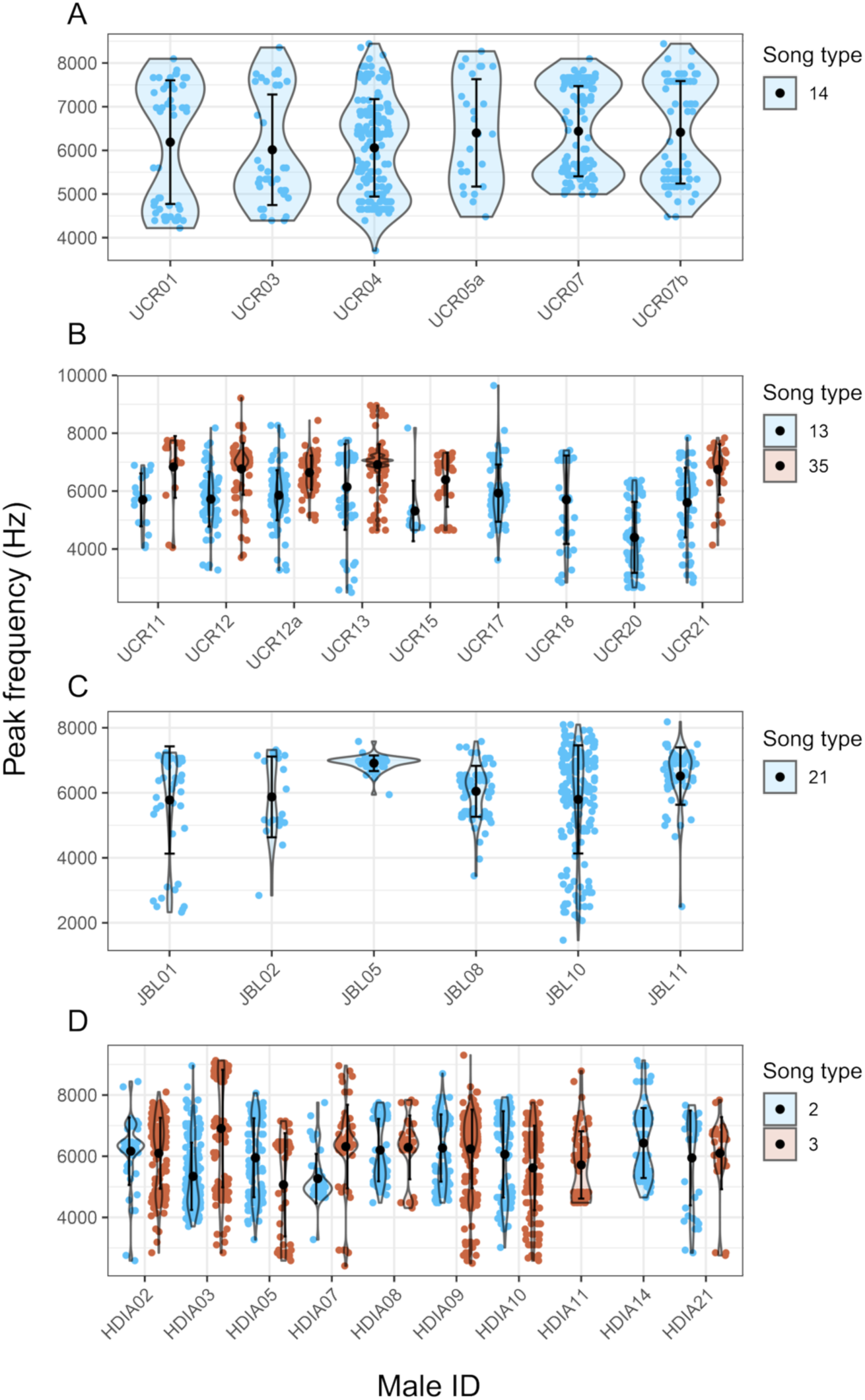
Temporal variance in frequency of maximum amplitude in common song types from (A) Highly urban, (B) Medium urban, (C) Low urban, and (D) Rural populations. Central black dots represent mean peak frequency for each male within the 13-time frame, error bars represent the standard deviation, and colored points represent a measurement.

## DISCUSSION

We analyzed the temporal consistency of common song types in the White-eared Ground-Sparrow over a 13-year period and found significant variation in fine-scale spectrotemporal song characteristics, both within-and between-individuals. Within-individual variation was greater than between-individual variation for common song types from the Medium, Low urbanized, and Rural populations, whereas between-individual variation exceeded within-individual variation in the Highly urbanized population. Despite these population-specific differences, analyses across the entire study period revealed that within-individual variation generally exceeded between-individual variation. Thus, our prediction that song structure would exhibit greater within-individual variation than between-individual variation through time was supported by three populations (Medium, Low urbanized, and Rural).

Our results revealed substantial between-individual variation in song structure, particularly in the Highly urbanized population. Differences among individuals in behavioral traits are often attributed to variation in environmental conditions, learning experiences, developmental processes, or genetic factors (Dingemanse et al. 2010; Ghalambor et al. 2010). For example, variation in song tempo in Bengalese Finches (*Lonchura striata domestica*) has a heritable genetic basis, although its contribution depends on birds’ song-learning experience (Mets & Brainard 2018), while early developmental stress alters song complexity in male Song Sparrows (*Melospiza melodia*) (Schmidt et al. 2014). Similar mechanisms may explain the patterns observed in White-eared Ground-Sparrows. Because this species exhibits post-natal dispersal (Rodríguez-Bardía et al. 2022), males are likely to experience different environmental conditions before settling into breeding territories, where they subsequently learn the local song repertoire (Sandoval et al. 2014; Cueva et al. 2025). Furthermore, dispersing males originate from multiple breeding populations (Rodríguez-Bardía et al. 2022; Cueva et al. 2025), suggesting that both genetic differences and variation in developmental environments could influence morphology and neural development, ultimately leading individuals to produce the same song types with distinct spectrotemporal characteristics (Mets & Brainard 2018). The particularly high between-individual variation detected in the Highly urbanized population is consistent with this interpretation and may additionally reflect habitat characteristics, such as reduced vegetation cover and extensive impervious surfaces, which alter sound transmission properties and may influence song structure (Cueva et al. 2024). Together, these findings support the idea that variation in developmental history, genetic background, and environmental experience can shape song expression and contribute to the persistence of between-individual differences in song structure in White-eared Ground-sparrows.

Territory variation due to habitat structure may be driving higher between-individual variation in the spectro-temporal features of common song types from the Highly urbanized population, with the level of urbanization playing a critical role (Méndez et al. 2021; Cueva et al. 2024). White-eared Ground-Sparrow territories in this site are surrounded by anthropogenic structures and are more open compared to the less urbanized site (Juárez et al. 2020), which ultimately limits transmission of certain acoustic traits (Gall et al. 2012; Cueva et al. 2024; Grimes et al. 2024). Considering that territories in this site are not homogeneous, and some resemble their natural habitat, males could be adjusting their songs according to their territory’s song transmission characteristics. In turn, males emit the same song types with varying spectral features, thus causing higher variation between-individuals. Moreover, variation in habitat structure between territories of the same population may also determine how concealed males are from anthropogenic disturbances. In Collared Flycatchers (*Ficedulla albicollis*), there were among individual differences in singing males depending on their perceived predation risk due to their singing position (Jablonsky et al. 2022). Given this, there is a chance that between-individual variation in the Highly urbanized population was also caused by how proximate males are to humans, which could be viewed by White-eared Ground-Sparrows as potential predators. Nonetheless, it has been argued that ambient noise rather than habitat structure poses as a stronger selective pressure on song transmission (LaZerte et al. 2015; Villarreal et al. 2024), suggesting that in urban spaces, anthropogenic noise might be mediating differences in song production between individuals (Harding et al. 2019; Méndez et al. 2021).

Over time within-individual variation was larger at the Medium urbanized, Low urbanized, and Rural populations. Meanwhile, the Highly urbanized population showed very low within-individual variation over time. We expected to find similar levels of within-individual variation between the Highly and Medium urbanized populations, because both sites are separated by 800 m.. This difference may be caused by differences in habitat structure or in the soundscape (Joo et al. 2011; Juárez et al. 2021; Méndez et al. 2021). The Highly urbanized site has been demonstrated to have high levels of anthropogenic noise compared to the Medium urbanized one (Juarez et al. 2017; Méndez et al. 2021). This may be influencing White-eared Ground-Sparrow males to limit the range of variation of their solo songs to avoid masking (Figure S1) as in Common Blackbirds (*Turdus merula*) and Southern House Wrens (*Troglodytes musculus*), which tended to produce more high-frequency elements to increase their communication in noisy urban environments (Nemeth et al. 2013; Juárez et al. 2021). As well, territories at the Highly urbanized site are more open than those at the Medium urbanized site (Juárez et al. 2020), and perhaps males could be striving to emit their solo songs more consistently to ensure proper transmission, since their original habitats are dense thickets (Cueva et al. 2024). White-eared Ground-Sparrow males in the Highly urbanized may be responding to anthropogenically induced environmental pressures (Méndez et al. 2021; Cueva et al. 2024).

Our results indicate that the Medium urbanized and Rural populations had the most within-individual temporal variation in their common song types (T13 and T35; T3), followed by the Low urbanized in two song characteristics for both of its song types (T21 and T42), which suggests less temporal consistency (Wilson 2018). Within-individual temporal variations in songs can emerge as males age, a trade-off that may occur to prevent further physiological deterioration rather than allocating resources to maintain their singing performance (Hunt et al. 2004; Bonduriansky et al. 2008). Previous studies suggested that Great Tit (*Parus major*) males at advanced ages tended to produce less consistent songs, while males of intermediate ages were more consistent (Rivera-Gutiérrez et al. 2012). Furthermore, temporal variations in songs associated with senescence can also arise due to physical constraints of older syrinxes that may ultimately affect motor performance and thus, song traits (Linville 1996). If this is the case, White-eared Ground-Sparrow common song types from these three populations seem to vary through time because of aging in males. However, if within-individual variation increases with age in this species, it fails to explain why males from the Highly urbanized population showed higher levels of between-individual variation, considering that they have higher survival rates than their rural counterparts (Juárez et al. 2022).

We found that the temporal variation in fine spectrotemporal features of common song types in White-eared Ground-Sparrow depends on their location. Between-individual variance predominated in the Highly urbanized population, a Highly urbanized site, suggesting that individual life histories and specific territorial conditions drive divergent temporal song trajectories among males. In contrast, within-individual variation was more prevalent in less urbanized populations, indicating less temporal consistency but a population-level response to time. These patterns might suggest that high urbanization levels may act as a threshold at which population-level responses give way to individual behavioral strategies. Our study contributes to understanding how time affect song variation and divergence at population level.

## Supporting information

Data & R scripts

## ACKNOWLEDGEMENTS

We thank the members of the Laboratorio de Ecología Urbana y Comunicación Animal (LEUCA) for recording part of the collected data. We thank Montserrat Alvarado-Deckwart and José Pablo Marín for data curation. We thank Universidad de Costa Rica and Jardín Botánico Lankester for the access to the study sites. We thank Vicerrectoría de Investigación for the financial support under investigation grant numbers C2705 and C3025.

## DATA AVAILABILITY

The data and code are provided as Supplementary Material and is also freely available in our Figshare repository, at: https://doi.org/10.6084/m9.figshare.33397525.

