## Supplemental tables and figures for "Consistency and individual variation in common song types of an urban bird: a multi-population comparison over a decade"

**Table S1**. Differences between White-eared Ground-Sparrow males in fine structure features from song types that presented between and within-individual variance.

| Population | Song type | Interactions | Min. freq. *p adj* | Max. freq. *p* adj | Bandwidth *p adj* | Peak freq. *p adj* | Song duration *p adj* |
| --- | --- | --- | --- | --- | --- | --- | --- |
| Highly urbanized | T14 | UCR03-UCR01 | **0.041** | 0.157 | **0.016** | 0.981 | **<0.001** |
|  |  | UCR04-UCR01 | **<0.001** | 0.260 | **0.002** | 0.980 | 0.110 |
|  |  | UCR05a-UCR01 | **0.010** | **<0.001** | **<0.001** | 0.974 | **<0.001** |
|  |  | UCR07-UCR01 | **0.038** | **0.022** | **0.002** | 0.800 | 0.324 |
|  |  | UCR07b-UCR01 | **<0.001** | **<0.001** | **<0.001** | 0.884 | **<0.001** |
|  |  | UCR04-UCR03 | 0.962 | 0.929 | 0.998 | 1.000 | **<0.001** |
|  |  | UCR05a-UCR03 | 0.976 | **<0.001** | **<0.001** | 0.781 | **<0.001** |
|  |  | UCR07-UCR03 | **<0.001** | **<0.001** | **<0.001** | 0.382 | **<0.001** |
|  |  | UCR07b-UCR03 | **<0.001** | **<0.001** | **<0.001** | 0.496 | **<0.001** |
|  |  | UCR05a-UCR04 | 0.999 | **<0.001** | **<0.001** | 0.725 | **<0.001** |
|  |  | UCR07-UCR04 | **<0.001** | **<0.001** | **<0.001** | 0.083 | 0.997 |
|  |  | UCR07b-UCR04 | **<0.001** | **<0.001** | **<0.001** | 0.200 | **0.002** |
|  |  | UCR07-UCR05a | **<0.001** | **<0.001** | **<0.001** | 1.000 | **<0.001** |
|  |  | UCR07b-UCR05a | **<0.001** | 0.992 | 0.467 | 1.000 | 0.150 |
|  |  | UCR07b-UCR07 | **<0.001** | **<0.001** | **<0.001** | 1.000 | **0.002** |
|  | T18 | UCR04-UCR03 | 1.000 | 0.999 | 0.999 | **<0.001** | **<0.001** |
|  |  | UCR05-UCR03 | **<0.001** | **0.043** | 0.274 | **<0.001** | **<0.001** |
|  |  | UCR06-UCR03 | 1.000 | 0.999 | 1.000 | **<0.001** | **<0.001** |
|  |  | UCR07-UCR03 | **0.0046** | **<0.001** | **<0.001** | **<0.001** | **<0.001** |
|  |  | UCR07b-UCR03 | **<0.001** | **<0.001** | **<0.001** | **<0.001** | **<0.001** |
|  |  | UCR08-UCR03 | **0.004** | **<0.001** | **<0.001** | **<0.001** | **0.001** |
|  |  | UCR05-UCR04 | **<0.001** | **0.023** | 0.027 | 0.126 | 0.091 |
|  |  | UCR06-UCR04 | 1.000 | 0.999 | 0.999 | 0.425 | 0.138 |
|  |  | UCR07-UCR04 | **<0.001** | **<0.001** | **<0.001** | **<0.001** | 0.231 |
|  |  | UCR07b-UCR04 | **<0.001** | **<0.001** | **<0.001** | 0.107 | 0.962 |
|  |  | UCR08-UCR04 | **<0.001** | **<0.001** | **<0.001** | 0.187 | **<0.001** |
|  |  | UCR06-UCR05 | **<0.001** | 0.072 | 0.281 | 1.000 | 0.999 |
|  |  | UCR07-UCR05 | **<0.001** | 0.053 | **<0.001** | 0.377 | 0.978 |
|  |  | UCR07b-UCR05 | **<0.001** | **<0.001** | **<0.001** | 1.000 | 0.487 |
|  |  | UCR08-UCR05 | **<0.001** | **<0.001** | **<0.001** | 1.000 | **<0.001** |
|  |  | UCR07-UCR06 | **<0.001** | **<0.001** | **<0.001** | 0.395 | 0.964 |
|  |  | UCR07b-UCR06 | **<0.001** | **<0.001** | **<0.001** | 1.000 | 0.523 |
|  |  | UCR08-UCR06 | **<0.001** | **<0.001** | **<0.001** | 1.000 | **<0.001** |
|  |  | UCR07b-UCR07 | **<0.001** | **<0.001** | **<0.001** | 0.101 | 0.851 |
|  |  | UCR08-UCR07 | 0.966 | **<0.001** | **<0.001** | 0.661 | **<0.001** |
|  |  | UCR08-UCR07b | **<0.001** | **<0.001** | **<0.001** | 1.000 | **<0.001** |
| Medium urbanized | T13 | UCR12-UCR11 | 0.808 | 0.177 | 0.123 | 1.000 | 0.545 |
|  |  | UCR12a-UCR11 | 0.899 | 1.000 | 1.000 | 1.000 | 0.054 |
|  |  | UCR13-UCR11 | **0.007** | 0.365 | **0.034** | 0.854 | **<0.001** |
|  |  | UCR15-UCR11 | 0.901 | 0.788 | 0.635 | 0.995 | 0.951 |
|  |  | UCR17-UCR11 | 0.763 | 0.987 | 1.000 | 0.997 | 1.000 |
|  |  | UCR18-UCR11 | 1.000 | 1.000 | 1.000 | 1.000 | 0.995 |
|  |  | UCR20-UCR11 | **<0.001** | 0.088 | **0.001** | **<0.001** | 0.270 |
|  |  | UCR21-UCR11 | 0.151 | 0.532 | 1.000 | 1.000 | **<0.001** |
|  |  | UCR12a-UCR12 | **<0.001** | **0.001** | **<0.001** | 0.997 | 0.759 |

Min. freq. = Minimum frequency; Max. freq. = Maximum frequency; Peak freq. = Peak frequency; *p adj.* = adjusted p-value. Bold values represent significant interactions.

Table S1. Continued.

| Population | Song type | Interactions | Min. freq. *p adj* | Max. freq. *p* adj | Bandwidth *p adj* | Peak freq. *p adj* | Song duration *p adj* |
| --- | --- | --- | --- | --- | --- | --- | --- |
|  |  | UCR13-UCR12 | **0.028** | 1.000 | 1.000 | 0.404 | **<0.001** |
| Medium urbanized | T13 | UCR15-UCR12 | 0.110 | **0.003** | **<0.001** | 0.980 | 1.000 |
|  |  | UCR17-UCR12 | **<0.001** | 0.210 | **0.002** | 0.973 | 0.180 |
|  |  | UCR18-UCR12 | 0.130 | **0.031** | **0.006** | 1.000 | 0.896 |
|  |  | UCR20-UCR12 | **<0.001** | 1.000 | 0.451 | **<0.001** | 1.000 |
|  |  | UCR21-UCR12 | 0.730 | **<0.001** | **<0.001** | 1.000 | **0.002** |
|  |  | UCR13-UCR12a | **<0.001** | **0.013** | **<0.001** | 0.784 | **<0.001** |
|  |  | UCR15-UCR12a | 1.000 | 0.498 | 0.581 | 0.883 | 0.994 |
|  |  | UCR17-UCR12a | 1.000 | 0.910 | 0.989 | 1.000 | **<0.001** |
|  |  | UCR18-UCR12a | 0.957 | 1.000 | 1.000 | 1.000 | 0.097 |
|  |  | UCR20-UCR12a | **<0.001** | **<0.001** | **<0.001** | **<0.001** | 0.993 |
|  |  | UCR21-UCR12a | **<0.001** | **0.004** | 1.000 | 0.843 | 0.120 |
|  |  | UCR15-UCR13 | **<0.001** | **0.010** | **<0.001** | 0.445 | **0.018** |
|  |  | UCR17-UCR13 | **<0.001** | 0.560 | **<0.001** | 0.972 | **<0.001** |
|  |  | UCR18-UCR13 | **<0.001** | 0.123 | **<0.001** | 0.571 | **<0.001** |
|  |  | UCR20-UCR13 | 0.865 | 0.987 | 0.911 | **<0.001** | **<0.001** |
|  |  | UCR21-UCR13 | 0.754 | **<0.001** | **<0.001** | 0.092 | 0.095 |
|  |  | UCR17-UCR15 | 1.000 | 0.171 | 0.320 | 0.807 | 0.961 |
|  |  | UCR18-UCR15 | 0.961 | 0.604 | 0.522 | 0.990 | 1.000 |
|  |  | UCR20-UCR15 | **<0.001** | **0.001** | **<0.001** | 0.310 | 1.000 |
|  |  | UCR21-UCR15 | **<0.001** | 1.000 | 0.785 | 1.000 | 0.495 |
|  |  | UCR18-UCR17 | 0.829 | 0.973 | 1.000 | 0.984 | 0.996 |
|  |  | UCR20-UCR17 | **<0.001** | 0.074 | **<0.001** | **<0.001** | **0.031** |
|  |  | UCR21-UCR17 | **<0.001** | **<0.001** | 0.856 | 0.697 | **<0.001** |
|  |  | UCR20-UCR18 | **<0.001** | **0.010** | **<0.001** | **<0.001** | 0.564 |
|  |  | UCR21-UCR18 | **<0.001** | 0.094 | 0.994 | 1.000 | **<0.001** |
|  |  | UCR21-UCR20 | **0.035** | **<0.001** | **<0.001** | **<0.001** | **0.026** |
|  | T35 | UCR12-UCR11 | **<0.001** | 0.713 | 0.201 | 1.000 | 0.993 |
|  |  | UCR12a-UCR11 | **<0.001** | 0.978 | **<0.001** | 0.912 | 0.605 |
|  |  | UCR13-UCR11 | **<0.001** | **0.001** | **0.003** | 0.998 | **0.003** |
|  |  | UCR15-UCR11 | **<0.001** | 0.963 | 0.803 | 0.287 | **<0.001** |
|  |  | UCR21-UCR11 | **<0.001** | 0.176 | **0.002** | 0.999 | **<0.001** |
|  |  | UCR12a-UCR12 | **<0.001** | **0.005** | <0.001 | 0.880 | **0.005** |
|  |  | UCR13-UCR12 | **<0.001** | **<0.001** | 0.001 | 0.672 | **<0.001** |
|  |  | UCR15-UCR12 | **<0.001** | 0.982 | 0.999 | 0.101 | **<0.001** |
|  |  | UCR21-UCR12 | **<0.001** | **<0.001** | 0.005 | 1.000 | **<0.001** |
|  |  | UCR13-UCR12a | **<0.001** | **<0.001** | <0.001 | 0.052 | **<0.001** |
|  |  | UCR15-UCR12a | **<0.001** | 0.245 | <0.001 | 0.505 | **<0.001** |
|  |  | UCR21-UCR12a | **<0.001** | 0.154 | 0.461 | 0.984 | **0.001** |
|  |  | UCR15-UCR13 | **<0.001** | **<0.001** | 0.015 | **0.001** | 0.172 |
|  |  | UCR21-UCR13 | **<0.001** | **<0.001** | 0.814 | 0.859 | **<0.001** |
|  |  | UCR21-UCR15 | 1.000 | **0.002** | 0.014 | 0.360 | **<0.001** |
| Low urbanized | T21 | JBL05-JBL01 | 0.775 | **0.018** | **0.002** | **<0.001** | 0.358 |
|  |  | JBL08-JBL01 | 0.261 | 0.957 | 1.000 | 0.825 | 0.555 |
|  |  | JBL10-JBL01 | 0.998 | 0.764 | 0.768 | 1.000 | **0.042** |

Min. freq. = Minimum frequency; Max. freq. = Maximum frequency; Peak freq. = Peak frequency; *p adj.* = adjusted p-value. Bold values represent significant interactions.

Table S1. Continued.

| Population | Song type | Interactions | Min. freq. *p adj* | Max. freq. *p* adj | Bandwidth *p adj* | Peak freq. *p adj* | Song duration *p adj* |
| --- | --- | --- | --- | --- | --- | --- | --- |
| Low urbanized | T21 | JBL11-JBL01 | **<0.001** | **<0.001** | **<0.001** | **0.042** | **<0.001** |
|  |  | JBL08-JBL05 | **0.011** | 0.053 | **<0.001** | **0.008** | 0.984 |
|  |  | JBL10-JBL05 | 0.430 | 0.063 | **0.005** | **<0.001** | 0.989 |
|  |  | JBL11-JBL05 | **0.004** | **<0.001** | **<0.001** | 0.579 | **<0.001** |
|  |  | JBL10-JBL08 | 0.141 | 0.991 | 0.670 | 0.557 | 0.692 |
|  |  | JBL11-JBL08 | **<0.001** | **<0.001** | **<0.001** | 0.271 | **<0.001** |
|  |  | JBL11-JBL10 | **<0.001** | **<0.001** | **<0.001** | **0.002** | **<0.001** |
|  | T42 | JBL02-JBL01 | **<0.001** | **<0.001** | 0.667 | **<0.001** | **<0.001** |
|  |  | JBL06-JBL01 | 0.129 | 0.998 | 1.000 | 0.409 | **<0.001** |
|  |  | JBL08-JBL01 | 1.000 | **<0.001** | 0.850 | **0.010** | 0.055 |
|  |  | JBL10-JBL01 | 0.960 | 0.077 | 0.994 | 0.793 | 0.205 |
|  |  | JBL11-JBL01 | 0.084 | **0.033** | 0.587 | **0.003** | 0.878 |
|  |  | JBL06-JBL02 | **<0.001** | **<0.001** | 0.649 | **<0.001** | 0.977 |
|  |  | JBL08-JBL02 | **<0.001** | 0.982 | 0.073 | **<0.001** | 0.435 |
|  |  | JBL10-JBL02 | **<0.001** | **<0.001** | 0.120 | **<0.001** | **0.002** |
|  |  | JBL11-JBL02 | **0.014** | 0.400 | **0.043** | **0.009** | **<0.001** |
|  |  | JBL08-JBL06 | 0.122 | **<0.001** | 0.944 | 0.826 | 0.203 |
|  |  | JBL10-JBL06 | **0.004** | 0.4219 | 1.000 | 0.877 | **0.002** |
|  |  | JBL11-JBL06 | 1.000 | 0.139 | 0.742 | 0.406 | **<0.001** |
|  |  | JBL10-JBL08 | 0.975 | **0.006** | 0.947 | **0.048** | 0.824 |
|  |  | JBL11-JBL08 | 0.080 | 0.807 | 0.990 | 0.950 | **0.008** |
|  |  | JBL11-JBL10 | **0.003** | 0.758 | 0.711 | **0.013** | **0.030** |
| Rural | T2 | HDIA03-HDIA02 | **0.002** | **0.004** | **<0.001** | **<0.001** | 0.618 |
|  |  | HDIA05-HDIA02 | **<0.001** | **<0.001** | **<0.001** | 0.976 | **<0.001** |
|  |  | HDIA07-HDIA02 | **<0.001** | **<0.001** | **<0.001** | **0.010** | **<0.001** |
|  |  | HDIA08-HDIA02 | **<0.001** | **<0.001** | **<0.001** | 1.000 | **<0.001** |
|  |  | HDIA09-HDIA02 | **<0.001** | **<0.001** | **<0.001** | 1.000 | **<0.001** |
|  |  | HDIA10-HDIA02 | **<0.001** | **<0.001** | **<0.001** | 1.000 | **<0.001** |
|  |  | HDIA14-HDIA02 | **<0.001** | **<0.001** | **<0.001** | 0.935 | **<0.001** |
|  |  | HDIA21-HDIA02 | 0.587 | **<0.001** | **<0.001** | 0.996 | **<0.001** |
|  |  | HDIA05-HDIA03 | **<0.001** | **<0.001** | **<0.001** | **<0.001** | **<0.001** |
|  |  | HDIA07-HDIA03 | **<0.001** | **<0.001** | **<0.001** | 1.000 | **<0.001** |
|  |  | HDIA08-HDIA03 | 0.970 | **<0.001** | **<0.001** | **<0.001** | **<0.001** |
|  |  | HDIA09-HDIA03 | **<0.001** | **0.012** | **<0.001** | **<0.001** | **0.002** |
|  |  | HDIA10-HDIA03 | **<0.001** | **<0.001** | **<0.001** | **<0.001** | **<0.001** |
|  |  | HDIA14-HDIA03 | **<0.001** | **0.010** | **<0.001** | **<0.001** | **<0.001** |
|  |  | HDIA21-HDIA03 | 0.928 | **<0.001** | **<0.001** | 0.118 | **0.007** |
|  |  | HDIA07-HDIA05 | 0.947 | 0.704 | 0.990 | **0.025** | 0.083 |
|  |  | HDIA08-HDIA05 | **<0.001** | 1.000 | 0.784 | 0.856 | **<0.001** |
|  |  | HDIA09-HDIA05 | 0.998 | **<0.001** | **<0.001** | 0.470 | **<0.001** |
|  |  | HDIA10-HDIA05 | 0.785 | 0.997 | 0.918 | 0.998 | **<0.001** |
|  |  | HDIA14-HDIA05 | **<0.001** | **<0.001** | 1.000 | **0.044** | **0.032** |
|  |  | HDIA21-HDIA05 | **<0.001** | 1.000 | 0.489 | 1.000 | **<0.001** |
|  |  | HDIA08-HDIA07 | **<0.001** | 0.695 | 1.000 | **0.001** | 0.694 |
|  |  | HDIA09-HDIA07 | 0.747 | 0.409 | 0.214 | **<0.001** | **0.008** |
|  |  | HDIA10-HDIA07 | 1.000 | 0.374 | 0.645 | **0.007** | 0.125 |
|  |  | HDIA14-HDIA07 | **<0.001** | 0.457 | 0.979 | **<0.001** | 1.000 |
|  |  | HDIA21-HDIA07 | **<0.001** | 0.809 | 0.987 | 0.211 | 0.438 |
|  |  | HDIA09-HDIA08 | **0.016** | **<0.001** | 0.199 | 1.000 | 0.446 |

Min. freq. = Minimum frequency; Max. freq. = Maximum frequency; Peak freq. = Peak frequency; *p adj.* = adjusted p-value. Bold values represent significant interactions.

Table S1. Continued.

| Population | Song type | Interactions | Min. freq. *p adj* | Max. freq. *p* adj | Bandwidth *p adj* | Peak freq. *p adj* | Song duration *p adj* |
| --- | --- | --- | --- | --- | --- | --- | --- |
| Rural | T2 | HDIA10-HDIA08 | **<0.001** | 1.000 | 0.160 | 0.997 | 0.979 |
|  |  | HDIA14-HDIA08 | **<0.001** | **<0.001** | 0.735 | 0.937 | 0.119 |
|  |  | HDIA21-HDIA08 | 0.569 | 1.000 | 1.000 | 0.978 | 1.000 |
|  |  | HDIA10-HDIA09 | 0.421 | **<0.001** | **<0.001** | 0.940 | 0.959 |
|  |  | HDIA14-HDIA09 | **<0.001** | 1.000 | **<0.001** | 0.987 | **<0.001** |
|  |  | HDIA21-HDIA09 | **<0.001** | **0.003** | 0.948 | 0.888 | 0.991 |
|  |  | HDIA14-HDIA10 | **<0.001** | **<0.001** | 0.985 | 0.375 | **<0.001** |
|  |  | HDIA21-HDIA10 | **<0.001** | 1.000 | 0.092 | 1.000 | 1.000 |
|  |  | HDIA21-HDIA14 | **<0.001** | **0.004** | 0.444 | 0.451 | 0.079 |
|  | T3 | HDIA03-HDIA02 | 0.073 | **0.005** | **0.002** | **<0.001** | **<0.001** |
|  |  | HDIA05-HDIA02 | **<0.001** | **<0.001** | **<0.001** | **<0.001** | **<0.001** |
|  |  | HDIA07-HDIA02 | 1.000 | **<0.001** | **<0.001** | 0.979 | **0.026** |
|  |  | HDIA08-HDIA02 | **<0.001** | **0.002** | **<0.001** | 0.999 | 0.984 |
|  |  | HDIA09-HDIA02 | **<0.001** | 0.099 | **<0.001** | 0.986 | **0.020** |
|  |  | HDIA10-HDIA02 | **0.001** | **<0.001** | **<0.001** | 0.051 | **0.028** |
|  |  | HDIA11-HDIA02 | **<0.001** | **<0.001** | **<0.001** | 0.645 | 0.595 |
|  |  | HDIA21-HDIA02 | **<0.001** | **<0.001** | **<0.001** | 1.000 | 0.997 |
|  |  | HDIA05-HDIA03 | **<0.001** | **<0.001** | **<0.001** | **<0.001** | **<0.001** |
|  |  | HDIA07-HDIA03 | 0.350 | **<0.001** | **<0.001** | 0.152 | 0.953 |
|  |  | HDIA08-HDIA03 | **<0.001** | **<0.001** | **<0.001** | 0.427 | **<0.001** |
|  |  | HDIA09-HDIA03 | **<0.001** | **<0.001** | **<0.001** | **0.001** | **<0.001** |
|  |  | HDIA10-HDIA03 | **<0.001** | **<0.001** | **<0.001** | **<0.001** | 0.171 |
|  |  | HDIA11-HDIA03 | **<0.001** | **<0.001** | **<0.001** | **<0.001** | 0.194 |
|  |  | HDIA21-HDIA03 | **<0.001** | **<0.001** | **<0.001** | 0.072 | 0.128 |
|  |  | HDIA07-HDIA05 | **0.006** | 0.534 | 0.051 | **<0.001** | **<0.001** |
|  |  | HDIA08-HDIA05 | 0.986 | 0.481 | 0.971 | **0.007** | 0.396 |
|  |  | HDIA09-HDIA05 | 1.000 | **<0.001** | **0.001** | **<0.001** | 0.371 |
|  |  | HDIA10-HDIA05 | 0.791 | 0.352 | 0.287 | 0.325 | **<0.001** |
|  |  | HDIA11-HDIA05 | 1.000 | 0.923 | 0.964 | 0.296 | **<0.001** |
|  |  | HDIA21-HDIA05 | 1.000 | 0.872 | 0.970 | **0.032** | **0.004** |
|  |  | HDIA08-HDIA07 | **<0.001** | 1.000 | 0.821 | 1.000 | **0.043** |
|  |  | HDIA09-HDIA07 | **<0.001** | **0.001** | 1.000 | 1.000 | **<0.001** |
|  |  | HDIA10-HDIA07 | **0.050** | 1.000 | 0.905 | **0.021** | 0.994 |
|  |  | HDIA11-HDIA07 | **<0.001** | **0.007** | **<0.001** | 0.265 | 0.958 |
|  |  | HDIA21-HDIA07 | **0.022** | 1.000 | 0.765 | 1.000 | 0.787 |
|  |  | HDIA09-HDIA08 | 0.998 | 0.214 | 0.400 | 1.000 | 1.000 |
|  |  | HDIA10-HDIA08 | 0.186 | 1.000 | 0.998 | 0.262 | 0.096 |
|  |  | HDIA11-HDIA08 | 0.999 | **0.018** | 0.381 | 0.633 | 0.402 |
|  |  | HDIA21-HDIA08 | 0.981 | 1.000 | 1.000 | 1.000 | 0.895 |
|  |  | HDIA10-HDIA09 | **0.009** | **<0.001** | 0.120 | **<0.001** | **<0.001** |
|  |  | HDIA11-HDIA09 | 1.000 | **<0.001** | **<0.001** | 0.159 | **<0.001** |
|  |  | HDIA21-HDIA09 | 0.999 | **0.012** | 0.300 | 1.000 | 0.135 |
|  |  | HDIA11-HDIA10 | 0.271 | **<0.001** | **<0.001** | 1.000 | 1.000 |
|  |  | HDIA21-HDIA10 | 0.914 | 1.000 | 0.999 | 0.652 | 0.965 |
|  |  | HDIA21-HDIA11 | 1.000 | 0.112 | 0.342 | 0.930 | 1.000 |

Min. freq. = Minimum frequency; Max. freq. = Maximum frequency; Peak freq. = Peak frequency; *p adj.* = adjusted p-value. Bold values represent significant interactions.

**Table S2**. Results from Wilcoxon test.

| Song type | Pillai’s Trace | Approx. *F* | *df* | den *df* | *p* |
| --- | --- | --- | --- | --- | --- |
| Highly urbanized |  |  |  |  |  |
| T14 | 1.088 | 35.49 | 20 | 1900 | <0.001 |
| T18 | 1.459 | 57.27 | 24 | 2392 | <0.001 |
| Medium urbanized |  |  |  |  |  |
| T13 | 0.714 | 16.17 | 28 | 2084 | <0.001 |
| T35 | 1.102 | 42.3 | 16 | 1780 | <0.001 |
| Low urbanized |  |  |  |  |  |
| T21 | 0.515 | 13.61 | 16 | 1472 | <0.001 |
| T42 | 0.454 | 10.84 | 20 | 1692 | <0.001 |
| Rural |  |  |  |  |  |
| T2 | 0.898 | 27.34 | 36 | 3400 | <0.001 |
| T3 | 0.581 | 16.01 | 40 | 3764 | <0.001 |

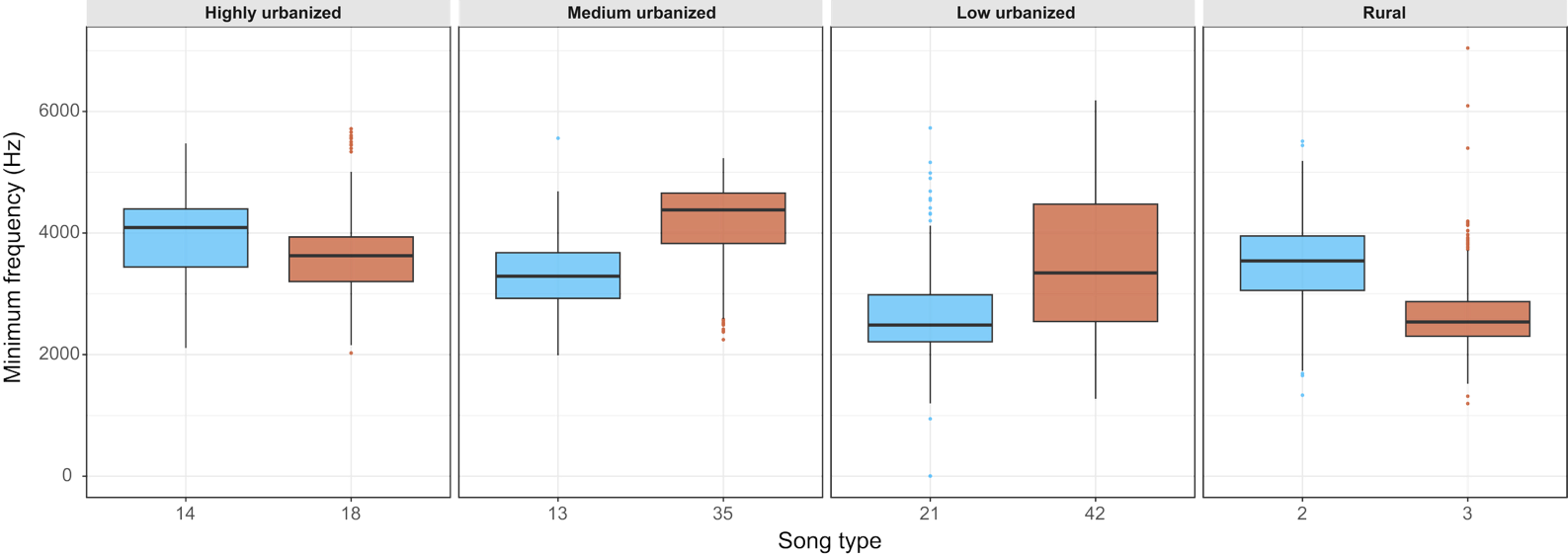

**Figure S1.** Descriptive statistics of minimum frequency for each song type from each population. Each box represents the interquartile range (25th–75th percentile), with the horizontal line indicating the median. Whiskers extend to 1.5 times the interquartile range, and points beyond the whiskers represent outliers.

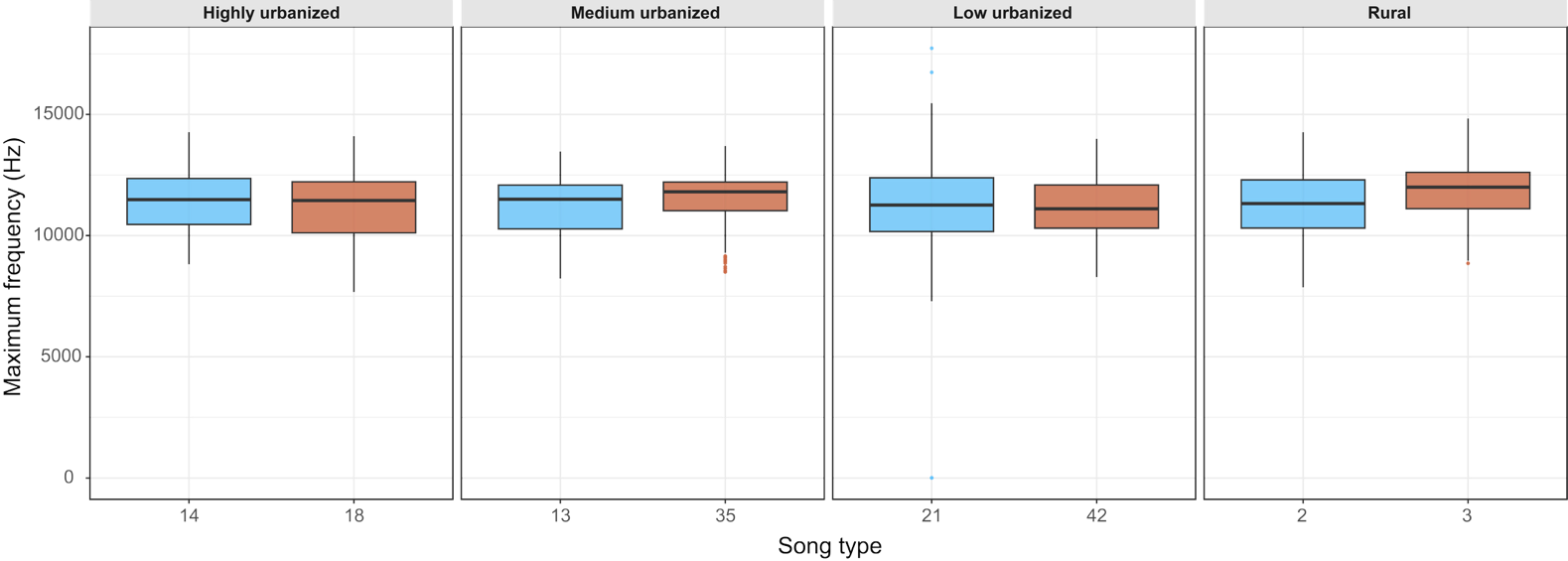

**Figure S2**. Descriptive statistics of maximum frequency for each song type from each population. Each box represents the interquartile range (25th–75th percentile), with the horizontal line indicating the median. Whiskers extend to 1.5 times the interquartile range, and points beyond the whiskers represent outliers.

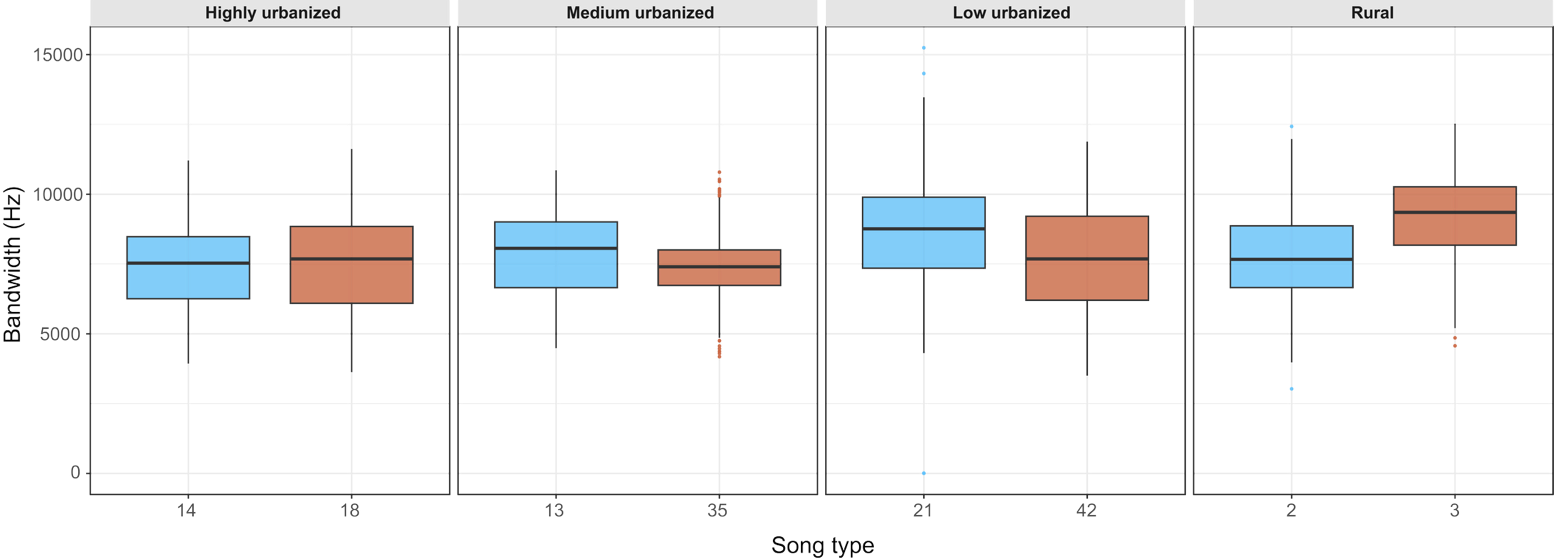

**Figure S3.** Descriptive statistics of frequency range for each song type from each population. Each box represents the interquartile range (25th–75th percentile), with the horizontal line indicating the median. Whiskers extend to 1.5 times the interquartile range, and points beyond the whiskers represent outliers.

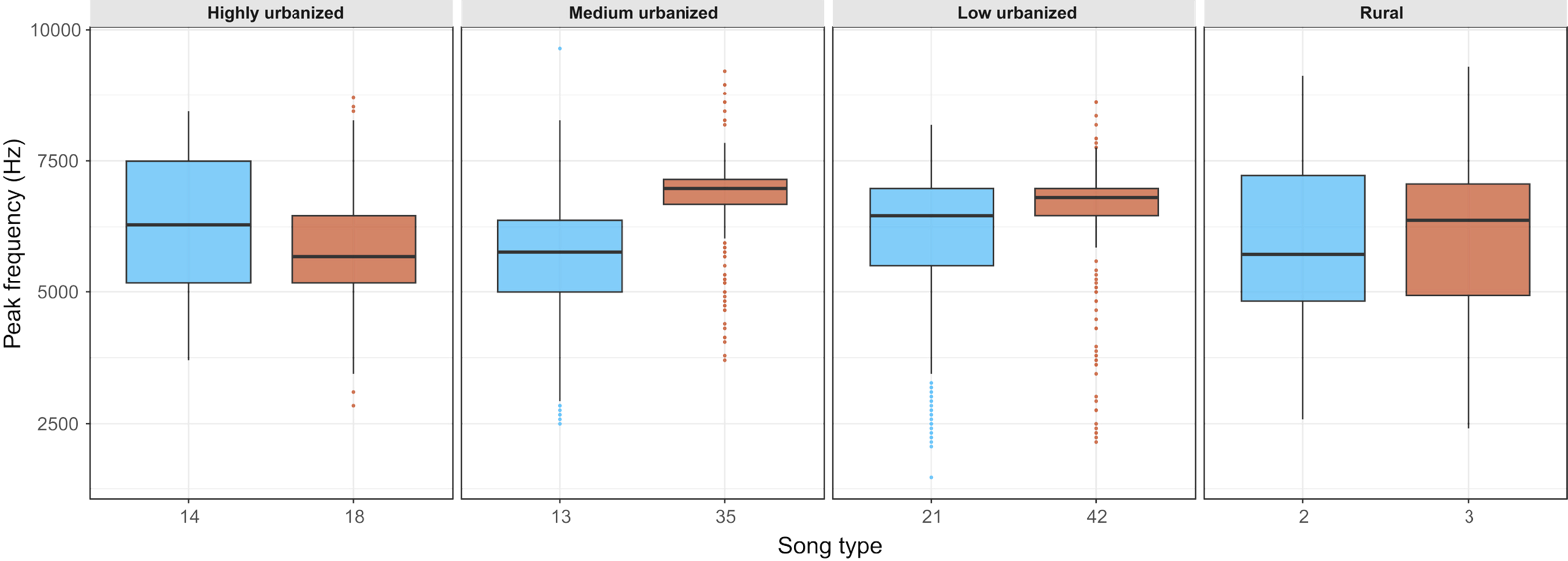

**Figure S4**. Descriptive statistics of frequency of maximum amplitude for each song type from each population. Each box represents the interquartile range (25th–75th percentile), with the horizontal line indicating the median. Whiskers extend to 1.5 times the interquartile range, and points beyond the whiskers represent outliers.

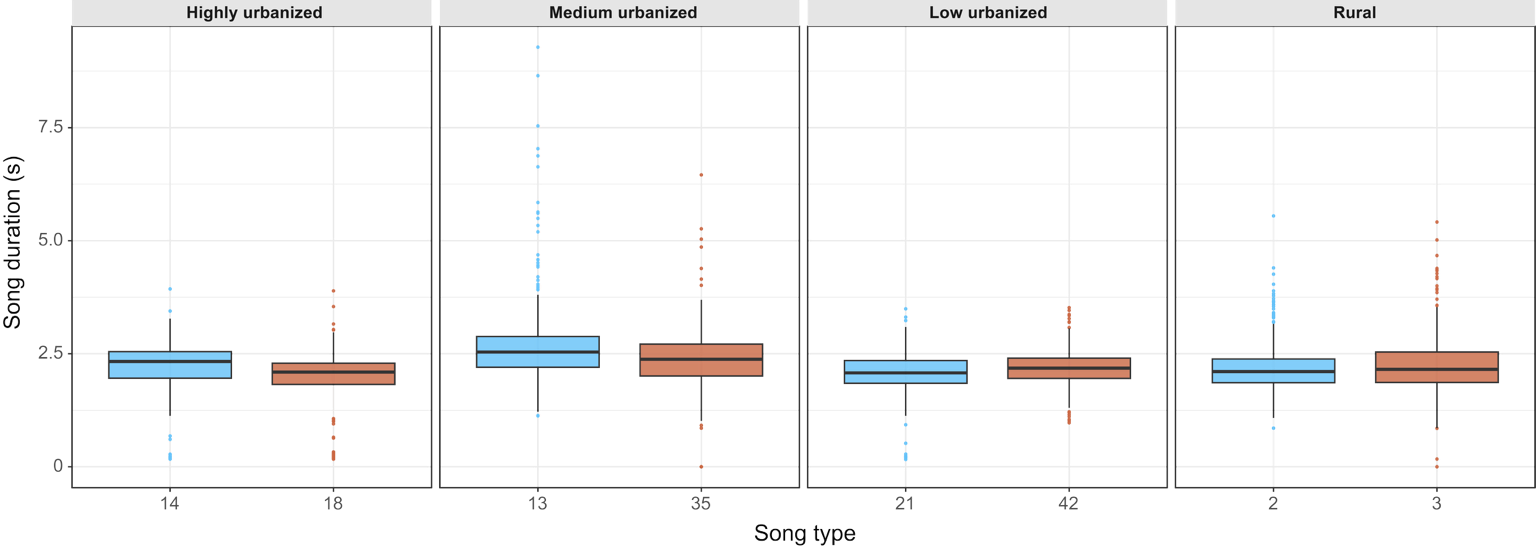

**Figure S5**. Descriptive statistics of song duration for each song type from each population. Each box represents the interquartile range (25th–75th percentile), with the horizontal line indicating the median. Whiskers extend to 1.5 times the interquartile range, and points beyond the whiskers represent outliers.
