## Supplementary material for "Consistency and individual variation in common song types of an urban bird: a multi-population comparison over a decade": Data & R scripts: Descriptive_Stats.Rmd

````
---
title: "Descriptive_Stats"
output: html_document
---

```{r setup, include=FALSE}
knitr::opts_chunk$set(echo = TRUE)
```

#1. Load libraries
```{r}
require(arm)
require(dplyr)
library(tidyverse)
```

#2. Reading Dataset
data: df_mleucotis.csv
```{r}
## ## SUBSTITUTE PATH TO DATA FOR "path"
data <- read.csv("path")
```


#3. Table A: Wilcoxon Test
```{r}

mis_variables <- c("Min_freq", "Max_freq", "Peak_freq", "Bandwidth", "Song_duration")
mis_etiquetas <- c("Minimum frequency (Hz)", "Maximum frequency (Hz)", 
                   "Peak frequency (Hz)", "Bandwidth (Hz)", "Song duration (s)")

comparaciones_completas <- lapply(
  c("Highly urbanized", "Medium urbanized", "Low urbanized", "Rural"),
  function(pob) {
    
    datos_pob <- df_mleucotis %>% filter(Population == pob)
    tipos     <- unique(datos_pob$Song_type)
    
    lapply(seq_along(mis_variables), function(idx) {
      
      nombre_var <- mis_variables[idx]
      etiqueta   <- mis_etiquetas[idx]
      
      grupo1 <- datos_pob %>% 
        filter(Song_type == tipos[1]) %>% 
        pull(!!sym(nombre_var))
      
      grupo2 <- datos_pob %>% 
        filter(Song_type == tipos[2]) %>% 
        pull(!!sym(nombre_var))
      
      grupo1 <- grupo1[!is.na(grupo1)]
      grupo2 <- grupo2[!is.na(grupo2)]
      
      test <- tryCatch(
        wilcox.test(grupo1, grupo2),
        error = function(e) NULL
      )
      
      if (!is.null(test)) {
        data.frame(
          Population   = pob,
          Variable     = etiqueta,
          Song_type_1  = tipos[1],
          Song_type_2  = tipos[2],
          Mean_T1      = round(mean(grupo1, na.rm = TRUE), 2),
          SD_T1        = round(sd(grupo1,   na.rm = TRUE), 2),
          Mean_T2      = round(mean(grupo2, na.rm = TRUE), 2),
          SD_T2        = round(sd(grupo2,   na.rm = TRUE), 2),
          W            = test$statistic,
          p_value      = round(test$p.value, 4),
          Significant  = ifelse(test$p.value < 0.05, "Yes", "No")
        )
      } else {
        data.frame(
          Population   = pob,
          Variable     = etiqueta,
          Song_type_1  = tipos[1],
          Song_type_2  = tipos[2],
          Mean_T1      = NA,
          SD_T1        = NA,
          Mean_T2      = NA,
          SD_T2        = NA,
          W            = NA,
          p_value      = NA,
          Significant  = NA
        )
      }
    }) %>% do.call(rbind, .)
  }) %>% do.call(rbind, .)

rownames(comparaciones_completas) <- NULL
print(comparaciones_completas)

```


#4. Table B: Acoustic description per song type and population 
```{r}

descriptives <- data %>%
  group_by(Population, Song_type) %>%
  summarise(
    n        = n(),
    
    Mean_MinFreq = round(mean(Min_freq, na.rm = TRUE), 1),
    SD_MinFreq    = round(sd(Min_freq,   na.rm = TRUE), 1),
    
    Mean_MaxFreq = round(mean(Max_freq, na.rm = TRUE), 1),
    SD_MaxFreq    = round(sd(Max_freq,   na.rm = TRUE), 1),
    
    Mean_Bandwidth = round(mean(Bandwidth, na.rm = TRUE), 1),
    SD_Bandwidth    = round(sd(Bandwidth,   na.rm = TRUE), 1),
    
    Mean_PeakFreq = round(mean(Peak_freq, na.rm = TRUE), 1),
    SD_PeakFreq    = round(sd(Peak_freq,   na.rm = TRUE), 1),
    
    Mean_SongDur = round(mean(Song_duration, na.rm = TRUE), 3),
    SD_SongDur    = round(sd(Song_duration,   na.rm = TRUE), 3),
    
    .groups = "drop"
  )
print(descriptives)

```
````
