## Supplementary material for "Consistency and individual variation in common song types of an urban bird: a multi-population comparison over a decade": Data & R scripts: Main_analysis.Rmd

````
---
title: "Song type consistency"
output: html_document
---

```{r setup, include=FALSE}
knitr::opts_chunk$set(echo = TRUE)
```

#1. Reading data
df_mleucotis.csv
```{r}
## ## SUBSTITUTE PATH TO DATA FOR "path"
data_frame <- read.csv("path")
```

#1. MANOVA
```{r}

## Highly urbanized ####

# Song type T14
t14_hu <- subset(data_frame, Song_type == 14 &  # Filtering data set for song type T14
                   Population == "Highly urbanized")

Y_T14 <- as.matrix(t14_hu[, c("Song_duration", "Min_freq", "Max_freq", "Peak_freq")]) # Generating matrix
man_t14 <- manova(Y_T14 ~ Unique.code, data = t14_hu)
summary(man_t14, test = "Pillai")

# Song type T18
t18_hu <- subset(data_frame, Song_type == 18 &  # Filtering data set for song type T18
                   Population == "Highly urbanized")

Y_T18 <- as.matrix(t18_hu[, c("Song_duration", "Min_freq", "Max_freq", "Peak_freq")]) # Generating matrix with acoustic features
man_t18 <- manova(Y_T18 ~ Unique.code, data = t18_hu)
summary(man_t18, test = "Pillai")

## Medium urbanized ####

# Song type T13
t13_mu <- subset(data_frame, Song_type == 13 &  # Filtering data set for song type T13
                   Population == "Medium urbanized") 

Y_T13 <- as.matrix(t13_mu[, c("Song_duration", "Min_freq", "Max_freq", "Peak_freq")])
man_t13 <- manova(Y_T13 ~ Unique.code, data = t13_mu)
summary(man_t13, test = "Pillai")

# Song type T35 
t35_mu <- subset(data_frame, Song_type == 35 &    # Filtering data set for song type T35
                   Population == "Medium urbanized")

Y_T35 <- as.matrix(t35_mu[, c("Song_duration", "Min_freq", "Max_freq", "Peak_freq")])
man_t35 <- manova(Y_T35 ~ Unique.code, data = t35_mu)
summary(man_t35, test = "Pillai")


## Low urbanized ####

# Song type T21
t21_lu <- subset(data_frame, Song_type == 21 &   # Filtering data set for song type T21
                   Population == "Low urbanized")

Y_T21 <- as.matrix(t21_lu[, c("Song_duration", "Min_freq", "Max_freq", "Peak_freq")])
man_t21 <- manova(Y_T21 ~ Unique.code, data = t21_lu)
summary(man_t21, test = "Pillai")

# Song type T42
t42_lu <- subset(data_frame, Song_type == 42 &    # Filtering data set for song type T42
                   Population == "Low urbanized")

Y_T42 <- as.matrix(t42_lu[, c("Song_duration", "Min_freq", "Max_freq", "Peak_freq")]) 
man_t42 <- manova(Y_T42 ~ Unique.code, data = t42_lu)
summary(man_t42, test = "Pillai")


## Rural ####

# Song type T2
t2_ru <- subset(data_frame, Song_type == 2 & 
                  Population == "Rural")   # Filtering data set for song type T2

Y_T2 <- as.matrix(t2_ru[, c("Song_duration", "Min_freq", "Max_freq", "Peak_freq")])
man_t2 <- manova(Y_T2 ~ Unique.code, data = t2_ru)
summary(man_t2, test = "Pillai")

# Song type T3
t3_ru <- subset(data_frame, Song_type == 3 &   # Filtering data set for song type T3
                  Population == "Rural")

Y_T3 <- as.matrix(t3_ru[, c("Song_duration", "Min_freq", "Max_freq", "Peak_freq")])
man_t3 <- manova(Y_T3 ~ Unique.code, data = t3_ru)
summary(man_t3, test = "Pillai")

```


#2. Function for LMM for each Song type and population
```{r}

# Main function
analysis_consistency <- function(data, name) {  # data = filtered data set; name = object name
  
  # Centering Year
  data <- data %>%
    mutate(Year_centered = Year - mean(Year, na.rm = TRUE))
  
  # Listing acoustic features
  variables <- c("Min_freq", "Max_freq", "Bandwidth", "Peak_freq", "Song_duration")
  
  results <- lapply(variables, function(var) {
    
    if (var != "Song_duration") data[[var]] <- data[[var]] / 1000  # Transforming to kHz for practicality for table
    
    # Linear Mixed Model formula
    formula <- as.formula(paste(var, "~ Year_centered + (1 | Unique.code)"))
    
    mod <- tryCatch(
      lmer(formula, data = data, REML = TRUE),
      error = function(e) NULL
    )
    
    if (is.null(mod)) {       # 
      return(data.frame(
        Dataset     = name,
        Variable    = var,
        Vind0       = NA,
        Ve0         = NA,
        VP          = NA,
        R           = NA,
        beta_year   = NA,
        p_year      = NA
      ))
    }
    
    # Extracting variances
    vc      <- as.data.frame(VarCorr(mod))
    Vind0   <- vc$vcov[vc$grp == "Unique.code"]  # Between-individual variance
    Ve0     <- vc$vcov[vc$grp == "Residual"]   # Within-individual variance
    VP      <- Vind0 + Ve0  # Phenotypic variance
    R       <- Vind0 / VP  # Repeatability
    
    # Extracting Year fixed effect 
    coefs   <- summary(mod)$coefficients
    beta    <- coefs["Year_centered", "Estimate"]
    p_val   <- coefs["Year_centered", "Pr(>|t|)"]
    
    data.frame(                    # Setting result table columns
      Dataset     = name,
      Variable    = var,
      Vind0       = round(Vind0, 4),
      Ve0         = round(Ve0, 4),
      VP          = round(VP, 4),
      R           = round(R, 4),
      beta_year   = round(beta, 4),
      p_year      = round(p_val, 5)
    )
  })
  
  do.call(rbind, results)
}

# Performing model for each population and song type

# Highly urbanized
res_hu_14 <- analysis_consistency(t14_hu, "Highly_Urb_T14")
res_hu_18 <- analysis_consistency(t18_hu, "Highly_Urb_T18")

# Medium urbanized
res_mu_13 <- analysis_consistency(t13_mu, "Medium_Urb_T13")
res_mu_35 <- analysis_consistency(t35_mu, "Medium_Urb_T35")

# Low urbanized
res_lu_21 <- analysis_consistency(t21_lu, "Low_Urb_T21")
res_lu_42 <- analysis_consistency(t42_lu, "Low_Urb_T42")

# Rural
res_ru_2 <- analysis_consistency(t2_ru, "Rural_T2")
res_ru_3 <- analysis_consistency(t3_ru, "Rural_T3")

# Combining results in table

table_results <- do.call(rbind, list(
  res_hu_14, res_hu_18,
  res_mu_13, res_mu_35,
  res_lu_21, res_lu_42,
  res_ru_2, res_ru_3
))

print(table_results)
```


#3. ANOVA for Highly urbanized population
```{r}
#### Song type T14 ####
#Filtering main data frame
t14_hu <- subset(data_frame, Song_type == 14 & 
                    Population == "Highly urbanized")

## ANOVA and Tukey tests

#Minimum frequency
f_lf <- aov(Min_freq ~ Unique.code, data = t14_hu)
summary(f_lf)
TukeyHSD(f_lf)  

#Maximum frequency
f_hf <- aov(Max_freq ~ Unique.code, data = t14_hu)
summary(f_hf)
TukeyHSD(f_hf)  

#Bandwidth
f_bw <- aov(Bandwidth ~ Unique.code, data = t14_hu)
summary(f_bw)
TukeyHSD(f_bw)

#Peak frequency
f_pf <- aov(Peak_freq ~ Unique.code, data = t14_hu)
summary(f_pf)
TukeyHSD(f_pf) 

#Song duration
f_dt <- aov(Song_duration ~ Unique.code, data = t14_hu)
summary(f_dt)
TukeyHSD(f_dt)

#### Song type T18 ####
t18_hu <- subset(data_frame, Song_type == 18 & 
                   Population == "Highly urbanized")

## ANOVA and Tukey test

#Minimum frequency
f_lf_18 <- aov(Min_freq ~ Unique.code, data = t18_hu)
summary(f_lf_18)
TukeyHSD(f_lf_18)  

#Maximum frequency
f_hf_18 <- aov(Max_freq ~ Unique.code, data = t18_hu)
summary(f_hf_18)
TukeyHSD(f_hf_18)

#Banddwidth
f_bw_18 <- aov(Bandwidth ~ Unique.code, data = t18_hu)
summary(f_bw_18)
TukeyHSD(f_bw_18) 

#Peak frequency
f_pf_18 <- aov(Peak_freq ~ Unique.code, data = t18_hu)
summary(f_pf_18)
TukeyHSD(f_pf_18)  

#Song duration
f_dt_18 <- aov(Song_duration ~ Unique.code, data = t18_hu)
summary(f_dt_18)
TukeyHSD(f_dt_18)

```

#4. ANOVA for Medium urbanized population
```{r}
#### Song type T13 ####
t13_mu <- subset(data_frame, Song_type == 13 & 
                   Population == "Medium urbanized") 

## ANOVA and Tukey test

#Minimum frequency
f_lf_13 <- aov(Min_freq ~ Unique.code, data = t13_mu)
summary(f_lf_13)
TukeyHSD(f_lf_13) 

#Maximum frequency
f_hf_13 <- aov(Max_freq ~ Unique.code, data = t13_mu)
summary(f_hf_13)
TukeyHSD(f_hf_13)  

#Bandwidth
f_bw_13 <- aov(Bandwidth ~ Unique.code, data = t13_mu)
summary(f_bw_13)
TukeyHSD(f_bw_13)

#Peak frequency
f_pf_13 <- aov(Peak_freq ~ Unique.code, data = t13_mu)
summary(f_pf_13)
TukeyHSD(f_pf_13)  

#Song duration
f_dt_13 <- aov(Song_duration ~ Unique.code, data = t13_mu)
summary(f_dt_13)
TukeyHSD(f_dt_13)


#### Song type T35 ####
t35_mu <- subset(data_frame, Song_type == 35 & 
                   Population == "Medium urbanized")

## ANOVA and Tukey test

#LF
f_lf_35 <- aov(Min_freq ~ Unique.code, data = t35_mu)
summary(f_lf_35)
TukeyHSD(f_lf_35) 

#Maximum frequency
f_hf_35 <- aov(Max_freq ~ Unique.code, data = t35_mu)
summary(f_hf_35)
TukeyHSD(f_hf_35)  

#Bandwidth
f_bw_35 <- aov(Bandwidth ~ Unique.code, data = t35_mu)
summary(f_bw_35)
TukeyHSD(f_bw_35)

#Peak frequency
f_pf_35 <- aov(Peak_freq ~ Unique.code, data = t35_mu)
summary(f_pf_35)
TukeyHSD(f_pf_35) 

#Song duration
f_dt_35 <- aov(Song_duration ~ Unique.code, data = t35_mu)
summary(f_dt_35)
TukeyHSD(f_dt_35)

```

#5. ANOVA for Low urbanized population 
```{r}
### Song type T21 ####
t21_lu <- subset(data_frame, Song_type == 21 & 
                   Population == "Low urbanized") 

## ANOVA and Tukey test

#Minimum frequency
f_lf_21 <- aov(Min_freq ~ Unique.code, data = t21_lu)
summary(f_lf_21)
TukeyHSD(f_lf_21)  

#Maximum frequency
f_hf_21 <- aov(Max_freq ~ Unique.code, data = t21_lu)
summary(f_hf_21)
TukeyHSD(f_hf_21)

#Bandwidth
f_bw_21 <- aov(Bandwidth ~ Unique.code, data = t21_lu)
summary(f_bw_21)
TukeyHSD(f_bw_21)

#Peak frequency
f_pf_21 <- aov(Peak_freq ~ Unique.code, data = t21_lu)
summary(f_pf_21)
TukeyHSD(f_pf_21)  

#Song duration
f_dt_21 <- aov(Song_duration ~ Unique.code, data = t21_lu)
summary(f_dt_21)
TukeyHSD(f_dt_21)

### Song type T42 #####
t42_lu <- subset(data_frame, Song_type == 42 & 
                   Population == "Low urbanized")

## ANOVA and Tukey test

#Minimum frequency
f_lf_42 <- aov(Min_freq ~ Unique.code, data = t42_lu)
summary(f_lf_42)
TukeyHSD(f_lf_42) 

#Maximum frequency
f_hf_42 <- aov(Max_freq ~ Unique.code, data = t42_lu)
summary(f_hf_42)
TukeyHSD(f_hf_42) 

#Bandwidth
f_bw_42 <- aov(Bandwidth ~ Unique.code, data = t42_lu)
summary(f_bw_42)
TukeyHSD(f_bw_42) 

#Peak frequency
f_pf_42 <- aov(Peak_freq ~ Unique.code, data = t42_lu)
summary(f_pf_42)
TukeyHSD(f_pf_42) 

#Song duration
f_dt_42 <- aov(Song_duration ~ Unique.code, data = t42_lu)
summary(f_dt_42)
TukeyHSD(f_dt_42)

```


#4. ANOVA for Rural population 
```{r}
### Song type T2 #####
t2_ru <- subset(data_frame, Song_type == 2 & 
                  Population == "Rural") 

## ANOVA and Tukey test

#Minimum frequency
f_lf_2 <- aov(Min_freq ~ Unique.code, data = t2_ru)
summary(f_lf_2)
TukeyHSD(f_lf_2)  

#Maximum frequency
f_hf_2 <- aov(Max_freq ~ Unique.code, data = t2_ru)
summary(f_hf_2)
TukeyHSD(f_hf_2)

#Bandwidth
f_bw_2 <- aov(Bandwidth ~ Unique.code, data = t2_ru)
summary(f_bw_2)
TukeyHSD(f_bw_2)   

#Peak frequency
f_pf_2 <- aov(Peak_freq ~ Unique.code, data = t2_ru)
summary(f_pf_2)
TukeyHSD(f_pf_2) 

#Song duration
f_dt_2 <- aov(Song_duration ~ Unique.code, data = t2_ru)
summary(f_dt_2)
TukeyHSD(f_dt_2)

### Song type T3 #####
t3_ru <- subset(data_frame, Song_type == 3 & 
                  Population == "Rural")

## ANOVA and Tukey test

#Minimum frequency
f_lf_3 <- aov(Min_freq ~ Unique.code, data = t3_ru)
summary(f_lf_3)
TukeyHSD(f_lf_3)  

#Maximum frequency
f_hf_3 <- aov(Max_freq ~ Unique.code, data = t3_ru)
summary(f_hf_3)
TukeyHSD(f_hf_3)

#Bandwidth
f_bw_3 <- aov(Bandwidth ~ Unique.code, data = t3_ru)
summary(f_bw_3)
TukeyHSD(f_bw_3) 

#Peak frequency
f_pf_3 <- aov(Peak_freq ~ Unique.code, data = t3_ru)
summary(f_pf_3)
TukeyHSD(f_pf_3)  

#Song duration
f_dt_3 <- aov(Song_duration ~ Unique.code, data = t3_ru)
summary(f_dt_3)
TukeyHSD(f_dt_3)

```
````
