## Supplementary material for "Consistency and individual variation in common song types of an urban bird: a multi-population comparison over a decade": Data & R scripts: ReadMe.docx

Purpose

This file explains how to use the attached data and code to conduct the analyses presented in the Supplemental Materials.

Data

The file df_mleucotis.csv is the base data of all the analysis conducted in the manuscript. It contains the following columns:

- **Recording.Code**: Name of recording; composed of the recorded individual ID, date, and recording day
- **Species**: Study species
- **Date**: Date of recording
- **Day**: Recording day.
- **Year**: Recording year.
- **Population**: Study population within the urban gradient.
- **Unique.code**: Male bird ID
- **Code**: Alternative male ID
- **Song_type**: Measured common song type within the recording
- **Start.time**: Time in seconds which song started within the recording
- **End.time:** Time in seconds in which song ended within the recording
- **Song_duration:** Song type length; calculated by End.time – Start.time
- **Min_freq:** Minimum frequency (Hz) of song type
- **Max_freq:** Maximum frequency (Hz) of song type
- **Peak_freq:** Peak frequency (Hz) of song type
- **Bandwidth:** Bandwidth of song type; calculated by Max_freq – Min_freq

Code

Rmd files can be opened in R Studio running Markdown and contain code used in the manuscript. They can be run in any order.

- Descriptive_Stats: Contains code for descriptive statistics.
- Main_analysis: Contains code for the linear mixed models, MANOVA, and ANOVA.
- Visualization: Contains code for graphs included in the manuscript and supplemental material.
