## Supplementary material for "Consistency and individual variation in common song types of an urban bird: a multi-population comparison over a decade": Data & R scripts: Visualization.Rmd

````
---
title: "Visualization"
output: html_document
---

```{r setup, include=FALSE}
knitr::opts_chunk$set(echo = TRUE)
```

## Loading packages
```{r}
library(ggplot2)
library(gridExtra)
```

data: df_mleucotis.csv
```{r}
## ## SUBSTITUTE PATH TO DATA FOR "path"
data_set <- read.csv("path")
```

#1. Generating main plots for each song type from each population for Figs. 3-7
```{r}

### Highly urbanized: Song types T14 y T18 ####
graph_hu <- data_set %>%      # Filtering data set for Highly urbanized 
  filter(Population == "Highly urbanized" & 
           Song_type %in% c("14", "18")) %>% 
  mutate(Song_type = as.factor(Song_type))


colors_hu <- c("14" = "#63C1F8", "18" = "#C96642") # Assigning colors to song types

# Generating individual figures for each measurement


## Minimum frequency

gglf_hu <- ggplot(graph_hu, aes(x = Unique.code, y = Min_freq, fill = Song_type)) +
  geom_point(aes(color = Song_type),
             position = position_jitterdodge(jitter.width = 0.5, dodge.width = 0.9),
             size = 1) +
  geom_violin(colour = "#00000099", 
              position = position_dodge(width = 0.9),
              alpha = 0.2,
              linewidth = 0.5) +
  stat_summary(fun = mean,
               geom = "point",
               position = position_dodge(width = 0.9),
               size = 1.5) +
  stat_summary(fun.data = mean_sdl,
               fun.args = list(mult = 1),
               geom = "errorbar",
               position = position_dodge(width = 1.0),
               width = 0.1,
               color = "black") +
  scale_fill_manual(values = colores_ucr) +
  scale_color_manual(values = colores_ucr) +
  labs(
    x = "Male ID",
    y = "Minimum frequency (Hz)",
    fill = "Song type",
    color = "Song type"
  ) +
  theme_bw() +
  theme(axis.text.x = element_text(angle = 45, hjust = 1))
gglf_hu


# Maximum frequency

gghf_hu <- ggplot(graph_hu, aes(x = Unique.code, y = Max_freq, 
                                fill = Song_type)) +
  geom_point(aes(color = Song_type),
             position = position_jitterdodge(jitter.width = 0.5, dodge.width = 0.9),
             size = 1) +
  geom_violin(colour = "#00000099", 
              position = position_dodge(width = 0.9),
              alpha = 0.2,
              linewidth = 0.5) +
  stat_summary(fun = mean,
               geom = "point",
               position = position_dodge(width = 0.9),
               size = 1.5) +
  stat_summary(fun.data = mean_sdl,
               fun.args = list(mult = 1),
               geom = "errorbar",
               position = position_dodge(width = 1.0),
               width = 0.1,
               color = "black") +
  scale_fill_manual(values = colores_ucr) +
  scale_color_manual(values = colores_ucr) +
  labs(
    x = "Male ID",
    y = "Maximum frequency (Hz)",
    fill = "Song type",
    color = "Song type"
  ) +
  theme_bw() +
  theme(axis.text.x = element_text(angle = 45, hjust = 1))
gghf_hu


# Bandwidth

ggbw_hu <- ggplot(graph_hu, aes(x = Unique.code, y = Bandwidth, fill = Song_type)) +
  geom_point(aes(color = Song_type),
             position = position_jitterdodge(jitter.width = 0.5, dodge.width = 0.9),
             size = 1)+
  geom_violin(colour = "#00000099", 
              position = position_dodge(width = 0.9),
              alpha = 0.2,
              linewidth = 0.5) +
  stat_summary(fun = mean,
               geom = "point",
               position = position_dodge(width = 0.9),
               size = 1.5) +
  stat_summary(fun.data = mean_sdl,
               fun.args = list(mult = 1),
               geom = "errorbar",
               position = position_dodge(width = 1.0),
               width = 0.1,
               color = "black") +
  scale_fill_manual(values = colores_ucr) +
  scale_color_manual(values = colores_ucr) +
  labs(
    x = "Male ID",
    y = "Bandwidth (Hz)",
    fill = "Song type",
    color = "Song type"
  ) +
  theme_bw() +
  theme(axis.text.x = element_text(angle = 45, hjust = 1))
ggbw_hu


#Peak frequency

ggpf_hu <- ggplot(graph_hu %>% filter(Song_type == "14"), 
                   aes(x = Unique.code, y = Peak_freq, fill = Song_type)) +
  geom_point(aes(color = Song_type),
             position = position_jitterdodge(jitter.width = 0.5, 
                                             dodge.width = 0.9),
             size = 1) +
  geom_violin(colour = "#00000099", 
              position = position_dodge(width = 0.9),
              alpha = 0.2,
              linewidth = 0.5) +
  stat_summary(fun = mean,
               geom = "point",
               position = position_dodge(width = 0.9),
               size = 1.5) +
  stat_summary(fun.data = mean_sdl,
               fun.args = list(mult = 1),
               geom = "errorbar",
               position = position_dodge(width = 1.0),
               width = 0.1,
               color = "black") +
  scale_fill_manual(values = colores_ucr) +
  scale_color_manual(values = colores_ucr) +
  labs(
    x = "Male ID",
    y = "Peak frequency (Hz)",
    fill = "Song type",
    color = "Song type"
  ) +
  theme_bw() +
  theme(axis.text.x = element_text(angle = 45, hjust = 1))
ggpf_hu


# Song duration

ggdt_hu <- ggplot(graph_hu, aes(x = Unique.code, y = Song_duration, fill = Song_type)) +
  geom_point(aes(color = Song_type),
             position = position_jitterdodge(jitter.width = 0.5, dodge.width = 0.9),
             size = 1) +
  geom_violin(colour = "#00000099", 
              position = position_dodge(width = 0.9),
              alpha = 0.2,
              linewidth = 0.5) +
  stat_summary(fun = mean,
               geom = "point",
               position = position_dodge(width = 0.9),
               size = 1.5) +
  stat_summary(fun.data = mean_sdl,
               fun.args = list(mult = 1),
               geom = "errorbar",
               position = position_dodge(width = 1.0),
               width = 0.1,
               color = "black") +
  scale_fill_manual(values = colores_ucr) +
  scale_color_manual(values = colores_ucr) +
  labs(
    x = "Male ID",
    y = "Song duration (s)",
    fill = "Song type",
    color = "Song type"
  ) +
  theme_bw() +
  theme(axis.text.x = element_text(angle = 45, hjust = 1))
ggdt_hu
               

```

```{r}
### Medium urbanized: Song types T13 y T35 ####
graf_mu <- data_set %>%      # Filtering data set for Medium urbanized
  filter(Population == "Medium urbanized" & 
           Song_type %in% c("13", "35")) %>%
  mutate(Song_type = as.factor(Song_type))


colores_dep <- c("13" = "#63C1F8", "35" = "#C96642") # Assigning colors to song types


# Generating individual figures for each measurement 

# Minimum frequency
gglf_mu <- ggplot(graf_mu, aes(x = Unique.code, y = Min_freq, 
                                 fill = Song_type)) +
  geom_point(aes(color = Song_type),
             position = position_jitterdodge(jitter.width = 0.5, dodge.width = 0.9),
             size = 1) +
  geom_violin(colour = "#00000099", 
              position = position_dodge(width = 0.9),
              alpha = 0.2,
              linewidth = 0.5) +
  stat_summary(fun = mean,
               geom = "point",
               position = position_dodge(width = 0.9),
               size = 1.5) +
  stat_summary(fun.data = mean_sdl,
               fun.args = list(mult = 1),
               geom = "errorbar",
               position = position_dodge(width = 1.0),
               width = 0.1,
               color = "black") +
  scale_fill_manual(values = colores_dep) +
  scale_color_manual(values = colores_dep) +
  labs(
    x = "Male ID",
    y = "Minimum frequency (Hz)",
    fill = "Song type",
    color = "Song type"
  ) +
  theme_bw() +
  theme(axis.text.x = element_text(angle = 45, hjust = 1))
gglf_mu


# Maximum frequency

gghf_mu <- ggplot(graf_mu, aes(x = Unique.code, y = Max_freq, fill = Song_type)) +
  geom_point(aes(color = Song_type),
             position = position_jitterdodge(jitter.width = 0.5, 
                                             dodge.width = 0.9),
             size = 1) +
  geom_violin(colour = "#00000099", 
              position = position_dodge(width = 0.9),
              alpha = 0.2,
              linewidth = 0.5) +
  stat_summary(fun = mean,
               geom = "point",
               position = position_dodge(width = 0.9),
               size = 1.5) +
  stat_summary(fun.data = mean_sdl,
               fun.args = list(mult = 1),
               geom = "errorbar",
               position = position_dodge(width = 1.0),
               width = 0.1,
               color = "black") +
  scale_fill_manual(values = colores_dep) +
  scale_color_manual(values = colores_dep) +
  labs(
    x = "Male ID",
    y = "Maximum frequency (Hz)",
    fill = "Song type",
    color = "Song type"
  ) +
  theme_bw() +
  theme(axis.text.x = element_text(angle = 45, hjust = 1))
gghf_mu


# Bandwidth

ggbw_mu <- ggplot(graf_mu, aes(x = Unique.code, y = Bandwidth, fill = Song_type)) +
  geom_point(aes(color = Song_type),
             position = position_jitterdodge(jitter.width = 0.5, dodge.width = 0.9),
             size = 1) +
  geom_violin(colour = "#00000099", 
              position = position_dodge(width = 0.9),
              alpha = 0.2,
              linewidth = 0.5) +
  stat_summary(fun = mean,
               geom = "point",
               position = position_dodge(width = 0.9),
               size = 1.5) +
  stat_summary(fun.data = mean_sdl,
               fun.args = list(mult = 1),
               geom = "errorbar",
               position = position_dodge(width = 1.0),
               width = 0.1,
               color = "black") +
  scale_fill_manual(values = colores_dep) +
  scale_color_manual(values = colores_dep) +
  labs(
    x = "Male ID",
    y = "Bandwidth (Hz)",
    fill = "Song type",
    color = "Song type"
  ) +
  theme_bw() +
  theme(axis.text.x = element_text(angle = 45, hjust = 1))
ggbw_mu


# Peak frequency

ggpf_mu <- ggplot(graf_mu, aes(x = Unique.code, y = Peak_freq, 
                               fill = Song_type)) +
  geom_point(aes(color = Song_type),
             position = position_jitterdodge(jitter.width = 0.5, dodge.width = 0.9),
             size = 1) +
  geom_violin(colour = "#00000099", 
              position = position_dodge(width = 0.9),
              alpha = 0.2,
              linewidth = 0.5) +
  stat_summary(fun = mean,
               geom = "point",
               position = position_dodge(width = 0.9),
               size = 1.5) +
  stat_summary(fun.data = mean_sdl,
               fun.args = list(mult = 1),
               geom = "errorbar",
               position = position_dodge(width = 1.0),
               width = 0.1,
               color = "black") +
  scale_fill_manual(values = colores_dep) +
  scale_color_manual(values = colores_dep) +
  labs(
    x = "Male ID",
    y = "Peak frequency (Hz)",
    fill = "Song type",
    color = "Song type"
  ) +
  theme_bw() +
  theme(axis.text.x = element_text(angle = 45, hjust = 1))
ggpf_mu


# Song duration

ggdt_mu <- ggplot(graf_mu, aes(x = Unique.code, y = Song_duration, fill = Song_type)) +
  geom_point(aes(color = Song_type),
             position = position_jitterdodge(jitter.width = 0.5, dodge.width = 0.9),
             size = 1) +
  geom_violin(colour = "#00000099", 
              position = position_dodge(width = 0.9),
              alpha = 0.2,
              linewidth = 0.5) +
  stat_summary(fun = mean,
               geom = "point",
               position = position_dodge(width = 0.9),
               size = 1.5) +
  stat_summary(fun.data = mean_sdl,
               fun.args = list(mult = 1),
               geom = "errorbar",
               position = position_dodge(width = 1.0),
               width = 0.1,
               color = "black") +
  scale_fill_manual(values = colores_dep) +
  scale_color_manual(values = colores_dep) +
  labs(
    x = "Male ID",
    y = "Song duration (s)",
    fill = "Song type",
    color = "Song type"
  ) +
  theme_bw() +
  theme(axis.text.x = element_text(angle = 45, hjust = 1))
ggdt_mu

```

```{r}
### Low urbanized: Song types T21 y 42 ####
graph_lu <- data_set %>%        # Filtering data set for Low urbanized
  filter(Population == "Low urbanized" & 
           Song_type %in% c("21", "42")) %>%
  mutate(Song_type = as.factor(Song_type))


colores_jbl <- c("21" = "#63C1F8", "42" = "#C96642") # Assigning colors to song types


# Generating individual figures for each measurement 


# Minimum frequency

gglf_lu <- ggplot(graph_lu, aes(x = Unique.code, y = Min_freq, 
                                fill = Song_type)) +
  geom_point(aes(color = Song_type),
             position = position_jitterdodge(jitter.width = 0.5, dodge.width = 0.9),
             size = 1) +
  geom_violin(colour = "#00000099", 
              position = position_dodge(width = 0.9),
              alpha = 0.2,
              linewidth = 0.5) +
  stat_summary(fun = mean,
               geom = "point",
               position = position_dodge(width = 0.9),
               size = 1.5) +
  stat_summary(fun.data = mean_sdl,
               fun.args = list(mult = 1),
               geom = "errorbar",
               position = position_dodge(width = 1.0),
               width = 0.1,
               color = "black") +
  scale_fill_manual(values = colores_jbl) +
  scale_color_manual(values = colores_jbl) +
  labs(
    x = "Male ID",
    y = "Minimum frequency (Hz)",
    fill = "Song type",
    color = "Song type"
  ) +
  theme_bw() +
  theme(axis.text.x = element_text(angle = 45, hjust = 1))
gglf_jbl

   
# Bandwidth

ggbw_lu <- ggplot(graph_lu %>% filter(Song_type == "42"), 
                   aes(x = Unique.code, y = Peak_freq, fill = Song_type)) +
  geom_point(aes(color = Song_type),
             position = position_jitterdodge(jitter.width = 0.5, dodge.width = 0.9),
             size = 1) +
  geom_violin(colour = "#00000099", 
              position = position_dodge(width = 0.9),
              alpha = 0.2,
              linewidth = 0.5) +
  stat_summary(fun = mean,
               geom = "point",
               position = position_dodge(width = 0.9),
               size = 1.5) +
  stat_summary(fun.data = mean_sdl,
               fun.args = list(mult = 1),
               geom = "errorbar",
               position = position_dodge(width = 1.0),
               width = 0.1,
               color = "black") +
  scale_fill_manual(values = colores_jbl) +
  scale_color_manual(values = colores_jbl) +
  labs(
    x = "Male ID",
    y = "Bandwidth (Hz)",
    fill = "Song type",
    color = "Song type"
  ) +
  theme_bw() +
  theme(axis.text.x = element_text(angle = 45, hjust = 1))
ggbw_lu


# Peak frequency

ggpf_lu <- ggplot(graph_lu %>% filter(Song_type == "21"), 
                   aes(x = Unique.code, y = Peak_freq, fill = Song_type)) +
  geom_point(aes(color = Song_type),
             position = position_jitterdodge(jitter.width = 0.5, dodge.width = 0.9),
             size = 1) +
  geom_violin(colour = "#00000099", 
              position = position_dodge(width = 0.9),
              alpha = 0.2,
              linewidth = 0.5) +
  stat_summary(fun = mean,
               geom = "point",
               position = position_dodge(width = 0.9),
               size = 1.5) +
  stat_summary(fun.data = mean_sdl,
               fun.args = list(mult = 1),
               geom = "errorbar",
               position = position_dodge(width = 1.0),
               width = 0.1,
               color = "black") +
  scale_fill_manual(values = colores_jbl) +
  scale_color_manual(values = colores_jbl) +
  labs(
    x = "Male ID",
    y = "Peak frequency (Hz)",
    fill = "Song type",
    color = "Song type"
  ) +
  theme_bw() +
  theme(axis.text.x = element_text(angle = 45, hjust = 1))
ggpf_lu


```

```{r}
### Rural: Song types T2 and T3 ####
graph_ru <- data_set %>%      # Filtering data set for Rural
  filter(Population == "Rural" & Song_type %in% c("3", "2")) %>%
  mutate(Song_type = as.factor(Song_type))


colores_hda <- c("2" = "#63C1F8", "3" = "#C96642") # Assigning colors to song types


# Generating individual figures for each measurement 

# Minimum frequency

gglf_ru <- ggplot(graph_ru, aes(x = Unique.code, y = Min_freq, 
                                fill = Song_type)) +
  geom_point(aes(color = Song_type),
             position = position_jitterdodge(jitter.width = 0.5, dodge.width = 0.9),
             size = 1) +
  geom_violin(colour = "#00000099", 
              position = position_dodge(width = 0.9),
              alpha = 0.2,
              linewidth = 0.5) +
  stat_summary(fun = mean,
               geom = "point",
               position = position_dodge(width = 0.9),
               size = 1.5) +
  stat_summary(fun.data = mean_sdl,
               fun.args = list(mult = 1),
               geom = "errorbar",
               position = position_dodge(width = 1.0),
               width = 0.1,
               color = "black") +
  scale_fill_manual(values = colores_hda) +
  scale_color_manual(values = colores_hda) +
  labs(
    x = "Male ID",
    y = "Minimum frequency (Hz)",
    fill = "Song type",
    color = "Song type"
  ) +
  theme_bw() +
  theme(axis.text.x = element_text(angle = 45, hjust = 1))
gglf_ru


# Maximum frequency

gghf_ru <- ggplot(graph_ru %>% 
                    filter(Song_type == "3"), 
                   aes(x = Unique.code, y = Max_freq, fill = Song_type)) +
  geom_point(aes(color = Song_type),
             position = position_jitterdodge(jitter.width = 0.5, dodge.width = 0.9),
             alpha = 0.5,
             size = 1) +
  geom_violin(colour = "#00000099", 
              position = position_dodge(width = 0.9),
              alpha = 0.2,
              linewidth = 0.5) +
  stat_summary(fun = mean,
               geom = "point",
               position = position_dodge(width = 0.9),
               size = 1.5) +
  stat_summary(fun.data = mean_sdl,
               fun.args = list(mult = 1),
               geom = "errorbar",
               position = position_dodge(width = 1.0),
               width = 0.1,
               color = "black") +
  scale_fill_manual(values = colores_hda) +
  scale_color_manual(values = colores_hda) +
  labs(
    x = "Male ID",
    y = "Maximum frequency (Hz)",
    fill = "Song type",
    color = "Song type"
  ) +
  theme_bw() +
  theme(axis.text.x = element_text(angle = 45, hjust = 1))
gghf_ru


# Bandwidth

ggbw_ru <- ggplot(graph_ru %>% filter(Song_type == "3"), 
                   aes(x = Unique.code, y = Bandwidth, 
                       fill = Song_type)) +
  geom_point(aes(color = Song_type),
             position = position_jitterdodge(jitter.width = 0.5, dodge.width = 0.9),
             size = 1) +
  geom_violin(colour = "#00000099", 
              position = position_dodge(width = 0.9),
              alpha = 0.2,
              linewidth = 0.5) +
  stat_summary(fun = mean,
               geom = "point",
               position = position_dodge(width = 0.9),
               size = 1.5) +
  stat_summary(fun.data = mean_sdl,
               fun.args = list(mult = 1),
               geom = "errorbar",
               position = position_dodge(width = 1.0),
               width = 0.1,
               color = "black") +
  scale_fill_manual(values = colores_hda) +
  scale_color_manual(values = colores_hda) +
  labs(
    x = "Male ID",
    y = "Bandwidth (Hz)",
    fill = "Song type",
    color = "Song type"
  ) +
  theme_bw() +
  theme(axis.text.x = element_text(angle = 45, hjust = 1))
ggbw_ru


# Peak frequency

ggpf_ru <- ggplot(graph_ru, aes(x = Unique.code, y = Peak_freq, 
                                fill = Song_type)) +
  geom_point(aes(color = Song_type),
             position = position_jitterdodge(jitter.width = 0.5, dodge.width = 0.9),
             size = 1) +
  geom_violin(colour = "#00000099", 
              position = position_dodge(width = 0.9),
              alpha = 0.2,
              linewidth = 0.5) +
  stat_summary(fun = mean,
               geom = "point",
               position = position_dodge(width = 0.9),
               size = 1.5) +
  stat_summary(fun.data = mean_sdl,
               fun.args = list(mult = 1),
               geom = "errorbar",
               position = position_dodge(width = 1.0),
               width = 0.1,
               color = "black") +
  scale_fill_manual(values = colores_hda) +
  scale_color_manual(values = colores_hda) +
  labs(
    x = "Male ID",
    y = "Peak frequency (Hz)",
    fill = "Song type",
    color = "Song type"
  ) +
  theme_bw() +
  theme(axis.text.x = element_text(angle = 45, hjust = 1))
ggmf_hda


#Song duration

ggdt_ru <- ggplot(graph_ru %>% filter(Song_type == "3"), 
                   aes(x = Unique.code, y = Song_duration, 
                       fill = Song_type)) +
  geom_violin(colour = "#00000099", 
              position = position_dodge(width = 0.9),
              alpha = 0.2,
              linewidth = 0.5) +
  geom_point(aes(color = Song_type),
             position = position_jitterdodge(jitter.width = 0.5, dodge.width = 0.9),
             size = 1) +
  stat_summary(fun = mean,
               geom = "point",
               position = position_dodge(width = 0.9),
               size = 1.5) +
  stat_summary(fun.data = mean_sdl,
               fun.args = list(mult = 1),
               geom = "errorbar",
               position = position_dodge(width = 1.0),
               width = 0.1,
               color = "black") +
  scale_fill_manual(values = colores_hda) +
  scale_color_manual(values = colores_hda) +
  labs(
    x = "Male ID",
    y = "Song duration (s)",
    fill = "Song type",
    color = "Song type"
  ) +
  theme_bw() +
  theme(axis.text.x = element_text(angle = 45, hjust = 1))
ggdt_ru

```
## Arranging figures
```{r}
# Minimum frequency (Fig.3)
grid.arrange(gglf_hu, gglf_mu, gglf_lu, gglf_ru, ncol = 1, nrow = 4)


# Maximum frequency (Fig. 4)
grid.arrange(gghf_hu, gghf_mu, gghf_ru, ncol=1, nrow=3)


# Bandwidth (Fig. 5)
grid.arrange(ggbw_hu, ggbw_mu, ggbw_lu, ggbw_ru, ncol=1, nrow=4)


# Peak frequency (Fig. 6)
grid.arrange(ggpf_hu, ggpf_mu, ggpf_lu, ggpf_ru, ncol=1, nrow=4)


# Song duration (Fig. 7)
grid.arrange(ggdt_hu, ggdt_mu, ggdt_ru, ncol=1, nrow=3)

```


#2. Boxplots for Supplementary materials
```{r}
 
# Filtering for population and song type 
data_plot <- data_set %>%
  filter(
    (Population == "Highly urbanized" & Song_type %in% c("14", "18")) |
    (Population == "Medium urbanized" & Song_type %in% c("13", "35")) |
    (Population == "Low urbanized"    & Song_type %in% c("21", "42")) |
    (Population == "Rural"            & Song_type %in% c("2", "3"))
  )

# Setting population as factor
data_set$Population <- factor(data_set$Population,
                       levels = c("Highly urbanized",
                                  "Medium urbanized",
                                  "Low urbanized",
                                  "Rural"))


# Boxplot function
boxplot_var <- function(variable, label_y){

  ggplot(data_plot,
         aes(x = Song_type,
             y = .data[[variable]],
             fill = Song_type)) +

    geom_boxplot(alpha = 0.7, outlier.size = 0.8) +

    facet_wrap(~Population,
               scales = "free_x",
               nrow = 1) +

    scale_fill_manual(values = colors_type) +

    labs(x = "Song type",
         y = label_y) +

    theme_bw(base_size = 11) +
    theme(
      legend.position = "none",
      strip.background = element_rect(fill = "grey90", color = NA),
      strip.text = element_text(face = "bold", size = 9)
    )
}

# Generating the five figures los 5 paneles
p_lf <- boxplot_var("Min_freq", "Minimum frequency (Hz)") #Fig S1
p_lf
p_hf <- boxplot_var("Max_freq", "Maximum frequency (Hz)") #Fig S2
p_hf
p_bw <- boxplot_var("Bandwidth", "Bandwidth (Hz)") #Fig S3
p_bw
p_pf <- boxplot_var("Peak_freq", "Peak frequency (Hz)") #Fig S4
p_pf
p_dt <- boxplot_var("Song_duration", "Song duration (s)") #Fig S5
p_dt

```
````
